# Targeted ATAC-see (tATAC-see): A Visual Assay for Target-Specific Chromatin Profiling

**DOI:** 10.64898/2026.07.13.738302

**Authors:** Natalie J. Kirkland, Yihao Yang, Manuel A. Castro, Pura Muñoz-Cánoves, Zach A. Levine

## Abstract

We present targeted ATAC-see (tATAC-see), a visual genomics assay for site-specific chromatin profiling. By integrating the *in situ* visualization of ATAC-see with antibody-tethered tagmentation, tATAC-see captures chromatin states at defined protein-occupied domains with the simplicity of standard immunofluorescence. Modulating the spatial interaction radius of Tn5 via salt titration enables extended tagmentation of local chromatin environments near the target. We validated this imaging-based method by capturing the expanded chromatin neighborhoods of active euchromatin (H3K27ac, H3K4me3) alongside the dense structural lamina-associated domains (LADs) of lamin A/C and lamin B1. Using an HDAC inhibitor, we tested the assay’s sensitivity to detect dynamic structural remodeling. We not only tracked chromatin decompaction but also demonstrated the robust structural resistance of LADs. As a proof-of-principle biological application, we applied tATAC-see to models of replicative, chronological, and pathological (Hutchinson–Gilford progeria syndrome) aging to assess its ability to detect well-characterized peripheral heterochromatin and LAD remodeling. We observed divergent chromatin trajectories at the nuclear envelope, reflective of the lamins’ distinct roles. lamin B1 domains exhibit increased local accessibility consistent with age-associated heterochromatin erosion, while lamin A/C-associated domains physically detach from their scaffold. Ultimately, tATAC-see provides a robust, accessible platform for mechanistic and population-level studies to uncover spatial epigenome dynamics.

**Summary Statement:** tATAC-see is a tunable visual genomics assay that can capture target-specific chromatin remodeling across diverse biological contexts.

## Introduction

Understanding the intricate architecture and dynamics of the cell requires not only observing which molecules are present but also understanding how they are spatially organized and functionally engaged. Within the highly compartmentalized mammalian nucleus, gene regulation is driven by mesoscale organization. The physical coupling of chromatin to specific domains, such as histone modifications, pioneer transcription factor hubs, and rigid structural boundaries, ultimately governs cell fate, cell state, and transcriptional output.

While traditional sequencing assays, such as ATAC-seq (Buenrostro et al., 2013), have revolutionized our global understanding of the epigenome, they require cells to be lysed in bulk, which removes spatial context and masks how chromatin accessibility fluctuates across distinct cell populations. To bridge this gap, recent advances have enabled spatially resolved epigenomic profiling through specialized microfluidic barcoding and multiplexed *in situ* hybridization platforms (Deng et al., 2022; Lu et al., 2022; Zhang et al., 2023). Alongside spatial sequencing, newer imaging-based methods now provide *in situ* visualization of chromatin accessibility. Tools such as ATAC-see and 3D ATAC-PALM allow analysis of the intranuclear distribution of accessible chromatin and have recently been adapted into cost-effective epigenetic screening platforms (Chen et al., 2016; Ishii et al., 2024; Kirkland et al., 2026; Xie et al., 2020).

Despite these advances, a fundamental gap remains. Global accessibility mapping cannot easily distinguish whether an open region of DNA is associated with an active transcription factor, a euchromatic histone mark, or structural scaffolds such as the nuclear lamina. Conversely, while targeted sequencing methods like CUT&Tag have successfully tethered the Tn5 transposase to specific antibodies to map protein–DNA interactions (Kaya-Okur et al., 2019; Kaya-Okur et al., 2020), determining the interacting chromatin environment of these binding footprints typically requires cross-comparison with parallel epigenomics datasets. Thus, linking locus-specific protein binding to the local chromatin environment within a native, visual context could be advantageous.

A recent variation of CUT&Tag, known as CUTAC, demonstrated that modifying the salt concentration of the tagmentation buffer allows tethered Tn5 to access nearby accessible chromatin neighborhoods. When targeting transcription-associated markers such as H3K4me2/3, this method revealed transcription-coupled accessible regulatory sites in the native genome (Henikoff et al., 2020). Combining the spatial resolution of ATAC-see with the antibody-tethering and salt-titration principles of CUTAC, we developed targeted ATAC-see (tATAC-see) to directly visualize site-specific chromatin architecture. By modulating the ionic strength of the reaction buffer, the effective interaction radius of the tethered Tn5 enzyme is tuneable: high-salt conditions restrict the enzyme to the immediate binding footprints of the target protein, while low-salt conditions allow it to probe the broader spatial neighborhood of tagmentable chromatin.

Coupled with an automated analysis pipeline, we rigorously validated the spatial and genomic fidelity of the assay using super-resolution microscopy and targeted DNA sequencing, demonstrating the expanded accessible local environments of active euchromatin alongside dense structural footprints of lamina-associated domains (LADs). To test the sensitivity of tATAC-see, we tracked acute chromatin decondensation following pharmacological histone deacetylase (HDAC) inhibition, revealing target-specific remodeling dynamics. Finally, we used tATAC-see to highlight the deterioration of nuclear lamina tethered chromatin during *in vitro* replicative aging as well as in human physiological and pathological aging. We revealed a divergent structural trajectory at the nuclear periphery, whereby aging induces a reduction in lamin A/C tagmentation efficiency while increasing the local tagmentability of lamin B1-associated domains. We propose that this reflects the physical detachment of chromatin from lamin A/C domains alongside the structural relaxation and erosion of heterochromatin at lamin B1 sites. Ultimately, with the technical ease of standard immunofluorescence, tATAC-see provides a robust, visual omics tool for uncovering complex local environmental dynamics of the epigenome, across histone marks, transcription factors, and architectural nuclear proteins.

## Results

### tATAC-see maps the target-proximal chromatin local environment with a tunable spatial radius

To directly visualize and quantify the chromatin architecture of specific protein-occupied local environments, we developed targeted ATAC-see (Fig. 1A). This approach merges the *in situ* visualization of ATAC-see (Chen et al., 2016) with the antibody-tethered tagmentation principles of CUT&Tag (Kaya-Okur et al., 2019; Kaya-Okur et al., 2020), and the low-salt conditions of CUTAC (Henikoff et al., 2020). We directed protein A/G-conjugated Tn5 to a target of interest via primary and fluorescent secondary antibodies in lightly fixed adherent cells (Fig. 1A, Fig. S1A). Because Tn5 was pre-loaded with fluorophore-labeled mosaic end (ME) oligonucleotides, active tagmentation at 37°C resulted in the direct, *in situ* incorporation of fluorescent marks into tagmentable regions of DNA. Crucially, the spatial reach of the tagmentation is governed by the ionic strength of the reaction buffer. Under restrictive high-salt conditions (300 mM NaCl), Tn5 cutting and insertion were limited to the DNA immediately juxtaposed to the target, yielding the visual equivalent of a strict CUT&Tag footprint. Conversely, under permissive low-salt conditions (0 mM NaCl), the flexibility of the tethered Tn5 complex increased, allowing it to rebind and tagment the broader spatial neighborhood (Henikoff et al., 2020), thereby capturing the tagmentability unique to each target’s specific local environment. Following targeted fluorescent insertion, stringent washes were applied to minimize the retention of unintegrated oligonucleotides (Fig. 1A). Nuclei were then counterstained with Hoechst to enable automated, high-throughput single-cell segmentation and spatial quantification.

**Figure 1.**
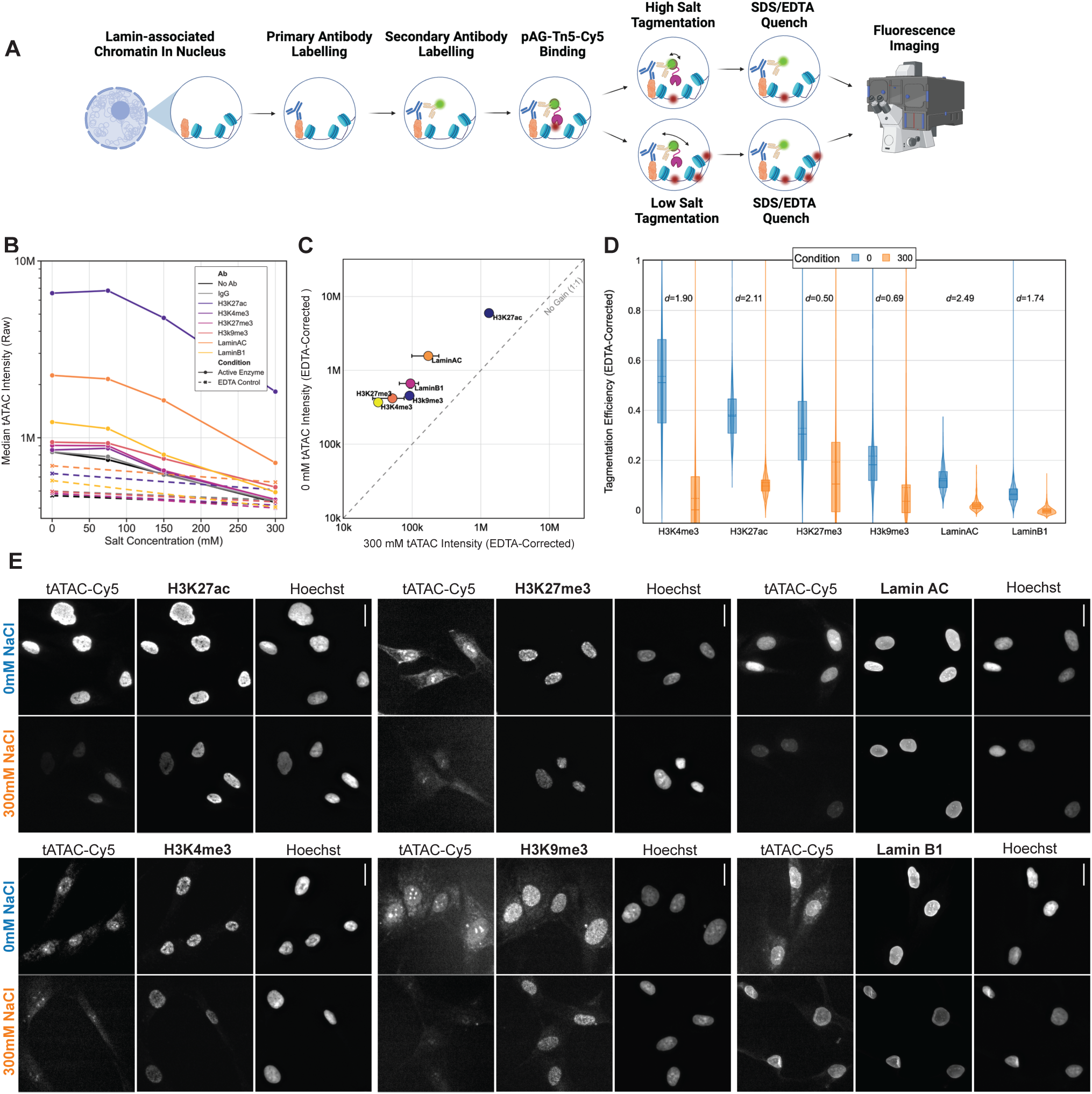
tATAC-see enables the spatial mapping and quantification of target-proximal chromatin local environments. (A) Schematic of the tATAC-see technique highlighting the primary and secondary antibody staining, pAG-Tn5 incubation, tagmentation and quenching steps prior to fluorescence imaging. (B) Line plot of raw median tATAC intensity across a salt titration gradient (0 to 300 mM NaCl) for diverse nuclear targets and negative controls (No Ab, IgG) applied to neonatal fibroblasts. Active enzyme reactions (solid lines) demonstrate a robust increase in accessibility at lower salt concentrations, whereas EDTA-inhibited controls (dashed lines) remain low across the gradient. (C) Scatter plot isolating the 300 mM standard footprint (x-axis) versus the0 mM expanded tagmentation (y-axis) using EDTA-subtracted tATAC intensity to control for background non-specific fluorescence and incomplete removal of non-integrated fluorescent adapters. Points resting above the dashed 1:1 "No Gain" line indicate local expansion at low salt. (D) Violin with box plots detailing the single-cell Tagmentation Efficiency (EDTA-Corrected), (tATAC intensity / Target intensity) - (EDTA tATAC intensity / Target intensity), for each target at 0 mM (blue) and 300 mM NaCl (orange), representing the chromatin tagged per unit of target. Boxes represent the interquartile range (IQR, 25th to 75th percentiles), with the solid line denoting the median. Dashed line is mean. Cohen’s *d* effect sizes (*d*) are annotated directly above each target pair. All 0 mM vs 300 mM comparisons yielded p < 0.0001 via two-sided Mann-Whitney U test. (B-D) Data shown are representative of a single 96-well plate experiment, which was performed twice for the full salt titration and more than three times for 0 and 300 mM pairs (*N* = 2; > 3 biological replicates). A minimum of *n* > 800 single nuclei were analyzed per condition. (E) Representative immunofluorescence images of active tATAC-see reactions at 0 mM and 300 mM NaCl. Columns display the tATAC-Cy5 signal alongside the specific target antibody footprint and Hoechst DNA staining. Scale bars = 20 µm.

To validate our method, we mapped the local chromatin environments at active histone marks (H3K4me3, H3K27ac), repressive heterochromatin (H3K9me3, H3K27me3), and the nuclear lamina (lamin A/C, lamin B1), alongside negative controls (no antibody, IgG), in primary human neonatal fibroblasts using a salt gradient of 300, 150, 75, and 0 mM NaCl. Because washes may not fully eliminate the physical retention of non-integrated oligonucleotides within the dense nuclear meshwork, we included EDTA control samples. By chelating the Mg^2+^ required for enzymatic catalysis, these controls established a baseline of non-specific retention, allowing us to assess and quantify background intensity. Tracking raw tATAC intensity from high-throughput widefield images with automated nuclear segmentation, we demonstrated that active Tn5 reactions expanded significantly at lower salt concentrations, while EDTA-inhibited controls remained consistently lower than matched experimental samples across the gradient (Fig. 1B). To evaluate assay robustness, we calculated the fold-change and signal-to-noise ratio (SNR) for each epigenetic and structural target. All targets exhibited SNRs greater than the non-specific IgG controls, confirming the spatial specificity of the initial tATAC-see intensities (Table S1).

Following EDTA subtraction, we further controlled for non-specific background fluorescence and incomplete removal of non-integrated fluorescent adapters (Fig. S1A, S2A, B) by comparing the restrictive high-salt state (300 mM NaCl) with the low-salt state (0 mM NaCl). This comparison also allowed us to define the localized chromatin footprint versus the expanded local environment, as a higher ratio of 0 mM to 300 mM indicates a more expansive tagmentable neighborhood of the target. H3K9me3 showed the lowest gain, while H3K4me3 showed the highest (Fig. 1C), reflecting the restrictive and permissive architectures of heterochromatin and euchromatin, respectively.

While all samples were exposed to 300 mM NaCl prior to tagmentation to prevent non-specific DNA-Tn5 binding, tagmentation at 37°C with 300 mM NaCl appeared to decrease target intensity for selected antibodies, including those against H3K27ac, lamin A/C, H3K27me3, and H3K4me3 (Fig. S2C). Consequently, differences between 0 and 300 mM tATAC-see intensities may be slightly exaggerated due to reduced Tn5 recruitment at 300 mM. As an additional metric to quantify local chromatin structure and to control for variations in target and antibody abundance (which directly affect the raw tATAC intensity), we calculated “tagmentation efficiency” as a measure of Tn5 activity (or DNA insertion events) per unit of target protein. To rigorously correct for non-specific background, we first normalized both the active and inactive (EDTA-corrected) tATAC signals by their respective target intensities, and then subtracted the median EDTA efficiency, to yield the final “tagmentation efficiency (EDTA-corrected)”. While the absolute values of this efficiency metric cannot be strictly compared across targets due to inherent differences in primary antibody affinity and fluorophore dynamics, their qualitative clustering aligns with established chromatin biology. At 0 mM NaCl, the euchromatic marks H3K4me3 and H3K27ac displayed higher DNA insertion events per target molecule, consistent with an open, nucleosome-depleted environment associated with active promoters, transcription, and enhancers (Fig. 1D). Conversely, lamin A/C and lamin B1 displayed the lowest absolute tagmentation efficiencies, likely reflecting the high abundance of these structural targets, the restricted surface area of the nuclear lamina-chromatin interface, and the inherently repressed chromatin local environments. The heterochromatic marks, H3K27me3 and H3K9me3, clustered together as an intermediate group, indicating a state more compacted than euchromatin but less structurally confined than the peripheral lamina. However, we note that nucleolar signals can be visualized for the heterochromatic marks, most prominently at 0 mM but also 300 mM (Fig. 1E and Fig. S1B). Profiling these specific repressive marks may be uniquely complicated by their localization to nucleolar-associated domains (NADs). Because the Tn5 enzyme kinetically prefers nucleosome-depleted DNA, tethering the enzyme to NADs likely results in the preferential tagmentation of the adjacent, highly accessible ribosomal DNA (rDNA) and euchromatin within the nucleolus, rather than the intended heterochromatin. Consequently, while they provide useful baseline metrics, these specific heterochromatic marks may be less optimal targets for evaluating true spatial expansion within this assay.

To overcome the limitations of absolute comparisons across targets, we used Cohen’s *d* to measure the effect size of the expansion between the restricted (300 mM) and unconstrained (0 mM) states (Fig. 1D). For euchromatic marks, the inherently open, nucleosome-depleted architecture allows for robust tagmentation even under restricted conditions, and extending the Tn5 reach yielded moderate spatial expansions relative to this high baseline (H3K27ac *d* = 2.11; H3K4me3 *d* = 1.90). For repressive heterochromatin marks, we observed small relative spatial expansions of the footprint when comparing the restrictive and unconstrained states (H3K27me3 *d* = 0.50; H3K9me3 *d* = 0.69). While lamin A/C and lamin B1 displayed the lowest tagmentation efficiencies, quantifying their expansion revealed striking differences. lamin A/C yielded a very large effect size (*d* = 2.49), reflective of a sharp transition from a tightly constrained footprint at the lamina-chromatin interface out to neighboring structural domains and potentially, the nucleoplasm. The lower relative efficiency and more restricted expansion of lamin B1 (*d =* 1.74) appears to align with its association with constitutive H3K9me3 (Steensel and Belmont, 2017), while lamin A/C is more frequently associated with dynamically regulated facultative heterochromatin and euchromatic marks (Gesson et al., 2016).

To verify that these observed increases were mathematically robust, we transformed the single-cell efficiencies into modified Z-scores. By standardizing the variance relative to the 300 mM baseline (Z=0), this metric confirmed a highly significant expansion of the tagmentation radius at 0 mM NaCl across all targeted domains with comparable target clustering observed with Cohen’s *d* (Fig. S2D).

Consistent with our quantitative metrics, direct visualization of the 20x widefield images confirmed that, for highly abundant targets (or targets with superior antibody binding kinetics), the 300 mM tATAC-see footprint colocalized with the target scaffold and subsequently expanded in intensity at 0 mM NaCl (Fig. 1E). Furthermore, visual evaluation of the corresponding EDTA-inhibited controls confirmed the loss of specific fluorescent signal across all targets, validating the technical specificity of the spatial assay (Fig. S1B).

Thus, this assay provides a robust visual and quantitative assessment of native chromatin states across diverse structural and epigenetic targets.

### High-resolution and genomic validation confirms that target-proximal local environments are precisely mapped using tATAC-see

Having established via high-throughput widefield imaging that modulating salt stringency drives a quantifiable expansion of the tATAC-see intensity (Fig. 1), we next sought to orthogonally validate the precise spatial fidelity and resolving power of these target-proximal local environments. To achieve this, we used Zeiss Airyscan super-resolution (SR) imaging on the active euchromatin marks H3K27ac and H3K4me3 in neonatal fibroblasts. Visualization confirmed that tATAC-see produced highly specific, target-anchored footprints (Fig. 2A, D). We then executed an automated pipeline to process 3D stack images, identify the central plane, mask the nucleus, and produce line scans, to allow unbiased assessment of quantitative colocalization and expansion metrics (Fig. S3A, D). Spatial line scans mapping the fluorescence intensities, globally scaled across the image set to enable relative comparison, further demonstrated specific tagmentation and elevated tATAC-see intensity at 0 mM NaCl relative to the target (Fig. 2B, E). To quantitatively assess colocalization, we calculated the Pearson correlation coefficient (PCC) and observed a high PCC for H3K27ac (0.8), which was unchanged by the salt concentration (Fig. S3B). In contrast, the highly punctate mark H3K4me3 exhibited significantly better spatial correlation at 0 mM NaCl (with an increase in PCC from 0.26 to 0.54; Fig. S3E), a reflection of the tagmentation signal smoothly broadening outward from the target anchor. To formally quantify the physical expansion of the tagmentation signal at the domain boundaries under low-salt conditions, we calculated the tagmentation to target ratio (relative tagmentation enrichment) across the target density gradient (Fig. S3C, F), which we used to define an edge broadening index at the low-intensity chromatin periphery (normalized target intensity of 0.1 to 0.3). For both euchromatic marks, the 0 mM condition demonstrated a highly significant physical expansion of the tagmentation signal beyond the strict target boundary as compared to the 300 mM baseline (Fig. 2C, F).

**Figure 2:**
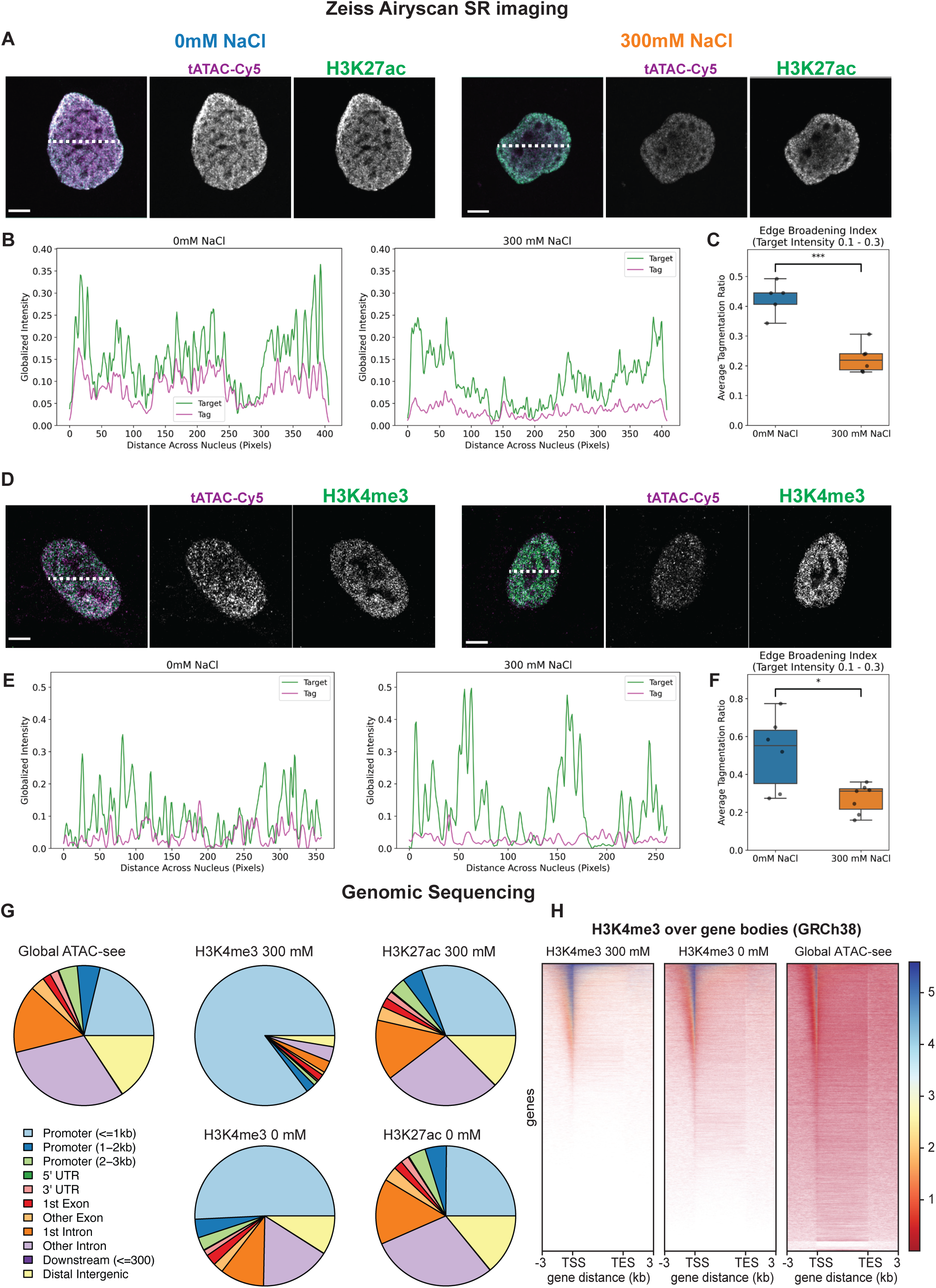
High-resolution mapping of tATAC-see demonstrates target-proximal accessible local environment of active histone modifications. (A-F) Airyscan SR imaging quantification for H3K27ac (A-C) and H3K4me3 (D-F) in neonatal fibroblasts. (A, D) Representative Airyscan SR cross-sectional images for H3K27ac (A) and H3K4me3 (D) tATAC-see at 0 and 300 mM NaCl, with dashed lines marking the regions used for the intensity profiles in (B, E). (B, E) Representative spatial line scans mapping the globally-scaled fluorescence intensities of the target (green) and tagmentation (magenta) channels at the cross-section of the nucleus marked in (A, D). The shared universal Y-axis demonstrates the suppression of tagmentation efficiency under high-salt (300 mM) conditions relative to the underlying target density. (C, F) Quantification of the Edge Broadening Index, defined as the average tagmentation to target ratio (plotted in Fig S3C, F) at the low-density chromatin periphery (Target Intensity 0.1 to 0.3). 0 mM NaCl shows a clear spatial expansion of the tagmentation intensity beyond the target compared to 300 mM NaCl. Independent Student’s t-tests were performed on n = 5; 6; 6; 7 single nuclei per condition for H3K27ac 0 and 300 mM, and H3K4me3 0 mM and 300 mM, respectively, (*p < 0.05, ***p < 0.001). (G) Genomic feature distribution of the peak sets (Global ATAC-see, H3K4me3 300 mM and 0 mM, H3K27ac 300 mM and 0 mM), annotated with ChIPseeker against hg38 genomic features. H3K4me3 300 mM was almost entirely promoter-proximal (≤1 kb) as expected, whereas H3K4me3 0 mM expands signal into interacting introns and distal intergenic space; H3K27ac (both conditions) and Global ATAC-see were broadly distributed across promoters, gene bodies, and distal intergenic regions. (H) Heatmaps of H3K4me3 300 mM, H3K4me3 0 mM, and Global ATAC-see signal over gene bodies (scale-regions, TSS−3 kb to TES+3 kb). Signal is concentrated at the TSS in all three conditions, presenting sharpest for 300 mM H3K4me3 and progressively broader for 0 mM and Global ATAC-see. Observations were confirmed for two independent cell lines.

To orthogonally validate these spatial expansions at the genomic level, we performed next-generation sequencing on amplified tATAC-see libraries generated from two independent cell lines. To accommodate sequencing requirements, we scaled the cell input to one well of a 12-well plate while strictly maintaining the identical in situ tATAC-see protocol. Following imaging, cells were reverse-crosslinked overnight prior to DNA extraction and library preparation. Peak calling confirmed the high biochemical specificity of the in situ tethering, as IgG controls yielded negligible peaks across cell lines (Fig. S3G). Consistent with our imaging data, shifting from 300 mM to 0 mM NaCl globally increased the total number of captured peaks for both marks (Fig. S3G). Annotation of genomic features revealed distinct architectural signatures driven by this spatial expansion. As expected, the 300 mM H3K4me3 footprint was almost entirely restricted to promoter-proximal regions (≤1 kb); however, extending the Tn5 radius at 0 mM allowed the signal to physically expand into interacting introns and intergenic spaces (Fig. 2G). Conversely, the H3K27ac signal was widely distributed across promoters, gene bodies, and distal intergenic regions regardless of salt stringency, reflecting its native distribution, and comparability to the Global ATAC-see library (equivalent to ATAC-seq) (Kirkland et al., 2026). UpSet plots further confirmed domain-specific architecture, demonstrating that H3K4me3 occupies a highly restricted fraction of the global accessible chromatin, whereas H3K27ac broadly co-occupies it (Fig. S3I, J). Signal distribution heatmaps over gene bodies further corroborated the specificity of H3K4me3 tATAC-see as the 300 mM H3K4me3 condition produced the sharpest focal peaks directly at the transcription start site (TSS), which progressively broadened at 0 mM but maintained localization versus the wider global ATAC-see landscape (Fig. 2H). For instance, at the *COL1A1* gene locus, both 0 mM and 300 mM tATACsee H3K4me3 signals were more enriched in the promoter as compared to the global ATACsee results. However, 0 mM yielded additional peaks at upstream and downstream enhancers (Fig. S3L). We next tested whether the expansion of H3K4me3 signal was restricted to interacting local chromatin across the genome. A genomic distance analysis demonstrated that the newly gained 0 mM H3K4me3 peaks were not random accessible background, but rather significantly clustered near the sharp 300 mM anchors compared to general non-promoter regions (Fig. S3K). Together, our analysis formally demonstrates that the low-salt signal expansion maps genuine, target-proximal chromatin neighborhoods.

Combined, the super-resolution and sequencing analyses establish an operational framework for interpreting epifluorescent tATAC-see readouts for euchromatin marks. Restrictive high-salt conditions tether Tn5 strictly to the immediately bound nucleosome, while low-salt conditions allow the enzyme to tagment physically-associated chromatin within the local 3D interaction hub, such as promoter–enhancer interactions (Fig. S3H).

### High-resolution and genomic mapping of tATAC-see captures continuous lamina-associated domains

We next sought to characterize the signal growth observed at the nuclear lamina in our initial widefield data, particularly given the inherently repressed nature of these domains (Fig. 1D). We applied the SR imaging and automated masking pipeline to lamin A/C and lamin B1 in neonatal fibroblasts (Fig. S4A, D). Direct visualization of the SR images clearly distinguished the antibody-targeted staining of the nuclear lamina and the fluorescently-tagmented proximal DNA. Because these channels represent distinct macromolecular structures at neighboring spatial coordinates, rather than perfectly overlapping entities, they exhibited visually distinct textures (Fig. 3A, D). Consequently, we observed fewer synchronous peaks in line scans (Fig. 3B, E) and moderate-to-low global colocalization scores (PCC ∼0.3–0.5) (Fig. S4B, E). Despite the distinct textures, the line scans confirmed that the tagmentation signal under permissive low-salt conditions was greater and was able to better track the target density (Fig. 3A, B, D, E), indicating that target-specific tagmentation had occurred. While the PCC trended higher at 0 mM, it was not statistically significant (Fig. S4B, E). Quantitatively profiling the relative tagmentation enrichment across the target density gradient revealed a significant spatial shift at 0 mM salt (Fig. 3C, F and Fig. S4C, F). The edge broadening index further confirmed that low salt drives physical expansion of the tagmentation signal beyond the strict lamina footprint for both lamin A/C and lamin B1.

**Figure 3:**
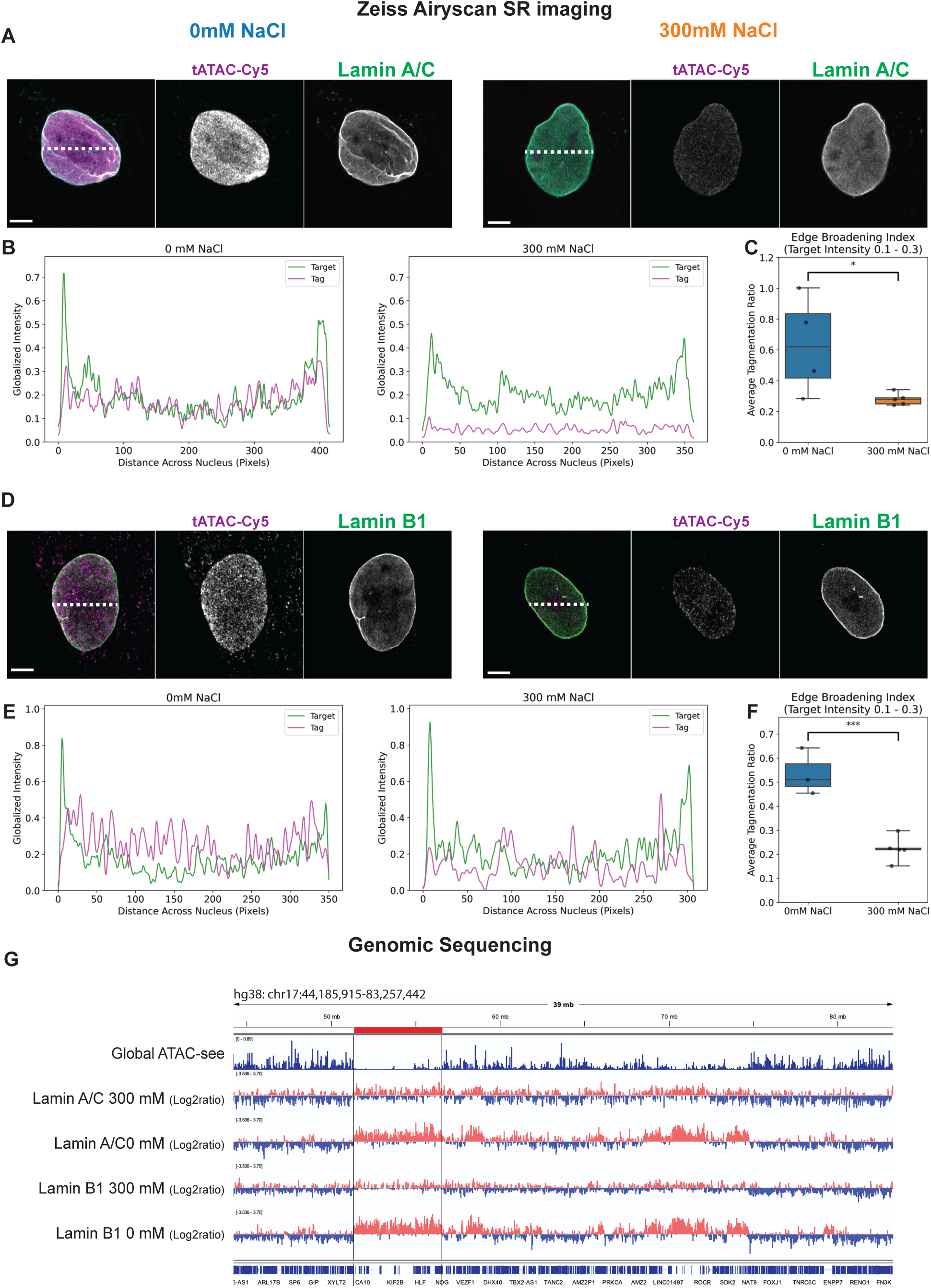
High-resolution mapping of tATAC-see demonstrates target-proximal tagmentable DNA at the nuclear lamina. (A-F) Airyscan SR imaging quantification for lamin A/C (A-C) and lamin B1 (D-F) in neonatal fibroblasts. (A, D) Representative Airyscan SR cross-sectional images for lamin A/C (A) and lamin B1 (D) tATAC-see at 0 and 300 mM NaCl, with dashed lines marking the regions used for the intensity profiles in (B, E). (B, E) Representative spatial line scans mapping the globally-scaled fluorescence intensities of the target (green) and tagmentation (magenta) channels at the cross-section of the nucleus marked in (A, D). The shared universal Y-axis demonstrates the suppression of tagmentation efficiency under high-salt (300 mM) conditions relative to the underlying target density. (C, F) Quantification of the Edge Broadening Index, defined as the average tagmentation to target ratio (plotted in Fig S4C, F) at the low-density chromatin periphery (Target Intensity 0.1 to 0.3). 0 mM NaCl shows a clear spatial expansion of the tagmentation intensity beyond the target compared to 300 mM NaCl. Independent Student’s t-tests were performed on *n* = 4; 5; 4; 5 single nuclei per condition for lamin A/C 0 and 300 mM, and lamin B1 0 mM and 300 mM, respectively, (*p < 0.05, ***p < 0.001). (G) IGV track view (hg38, chr17:44,185,915–83,257,442; ∼39 Mb) comparing Global ATAC-see accessibility with lamin A/C and lamin B1 nuclear-lamina association at 300 mM and 0 mM NaCl. lamin tracks represent the log2 ratio of each lamin sample over the 0 mM IgG control (red = Lamin-enriched, blue = Lamin-depleted). Observations were confirmed for two independent cell lines. The salt titration reshapes the lamina profile: at 300 mM, both lamin A/C and lamin B1 exhibit a punctate, discontinuous signal that sharply pinpoints individual lamina-contact anchors, whereas at 0 mM the signal expands into continuous blocks spanning entire Lamina-Associated Domains (e.g., the red highlighted region). This sharp-anchor to whole-domain broadening demonstrates that low ionic strength captures chromatin at the level of complete nuclear-architecture domains rather than discrete binding sites.

We therefore sought to investigate how this spatial expansion at the nuclear periphery translated to the genome. We first generated tATAC-see libraries as described for the euchromatin marks, but a lack of enrichment in accessible chromatin was detected by global ATAC-see (Fig. S4G) confirmed that the genomic architecture of these structural targets was highly distinct. We therefore processed the corresponding sequencing libraries using pipelines established for LADs and calculated the log2 ratio of the lamin tATAC-see signal over the 0 mM IgG control. This analysis revealed a distinct chromatin profile driven by the salt concentrations. At 300 mM NaCl, both lamin A/C and lamin B1 exhibited a punctate, discontinuous genomic profile, sharply pinpointing individual, discrete lamina-contact anchors. Conversely, shifting the assay to 0 mM NaCl caused these isolated anchors to expand into continuous, broad blocks that spanned entire LADs (Fig. 3G). Crucially, we did not observe erroneous tagmentation at globally accessible regions; in fact, visualization of the genomic tracks highlighted the expected anticorrelation, with the expanded lamin tATAC-see signal robustly marking gene-poor domains that are largely devoid of global ATAC-see accessibility. This sequencing data likely models the restrictive steric hindrance and limited surface area of the immediate lamina–chromatin interface, which the extended interaction radius at 0 mM can overcome to allow Tn5 to probe and tagment the surrounding LAD, which is largely heterochromatic.

Thus, our combined super-resolution and sequencing analyses establish the operational framework for using tATAC-see at structural elements. While high ionic strength restricts the signal to direct physical contact sites, low ionic strength effectively demarcates complete, higher-order nuclear-architecture domains without conflating their baseline tagmentability with euchromatic openness.

### tATAC-see detects dose-dependent chromatin remodeling following HDAC inhibition

We next evaluated the assay’s sensitivity to detect acute, dose-dependent changes in local chromatin architecture. We treated primary neonatal and 22-year-old dermal fibroblasts with trichostatin A (TSA), a potent, broad-spectrum histone deacetylase (HDAC) inhibitor that drives controlled chromatin decondensation. This provided a physiological model to assess whether tATAC-see can resolve subtle, progressive chromatin remodeling.

Previous CUTAC sequencing has shown that targeting H3K4me2/3 under low ionic conditions generates chromatin accessibility maps virtually indistinguishable from bulk ATAC-seq, as H3K4me3-tethered Tn5 at active promoters can tagment neighboring nucleosome-depleted regions (Henikoff et al., 2020). Since we have previously demonstrated that global ATAC-see detects TSA-driven chromatin opening (Kirkland et al., 2026), we hypothesized that targeting the euchromatic histone modifications H3K27ac and H3K4me3 would yield a measurable increase in tATAC-see intensity in low salt conditions. Indeed, a 4-hour treatment with 0, 125, and 250 nM TSA drove a dose-dependent increase in 0 mM NaCl tATAC intensity at both euchromatic targets and for two independent fibroblast lines (Fig. 4A, D; Fig. S5A, C, E, G). Because the total abundance of H3K27ac and H3K4me3 actively increased with TSA treatment as expected (Fig. S5B, D, F, H), the structural relaxation also translated into an increased 300 mM DNA footprint, equivalent to the standard CUT&Tag visualization (Fig. 4A, D and Fig. S5E).

**Figure 4:**
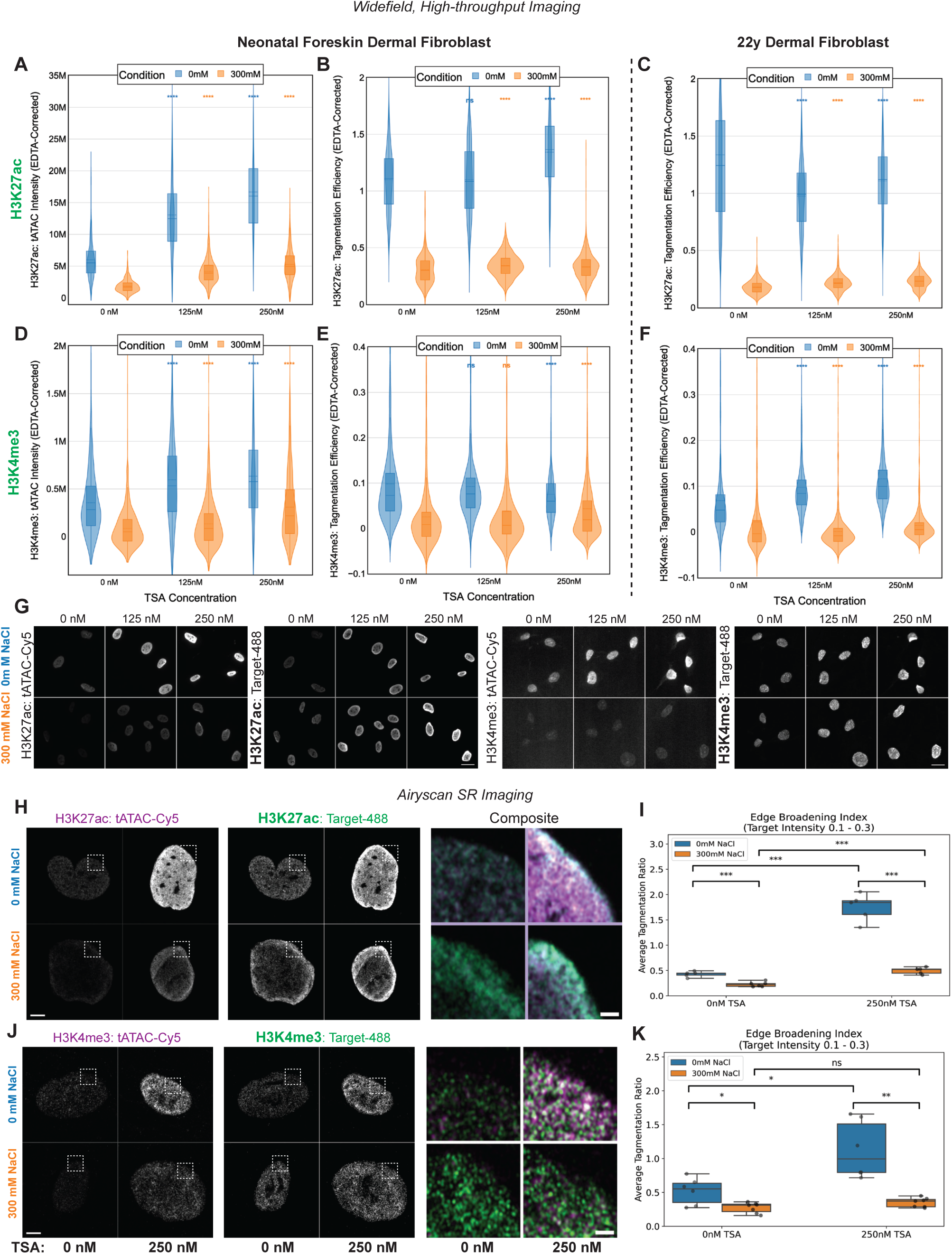
tATAC-see sensitively captures global chromatin remodeling following HDAC inhibition. (A–F) Quantitative evaluation of neonatal (A, B, D, E) and 22-year-old adult HDFs (C, F) subject to 4-hour treatment with 0, 125, or 250 nM Trichostatin A (TSA). Total tATAC-see intensity (EDTA-Corrected) (A, D) and normalized tagmentation efficiency (B, C, E, F) are plotted for H3K27ac (A-C) and H3K4me3 (D-F). (G) Representative extended depth of focus (EDF) projections from widefield imaging of the tagmentation signal (tATAC-Cy5) and the structural anchor (Target-488, H3K27ac or H3K4me3 primary antibody) across TSA concentrations and salt conditions (0 mM vs. 300 mM NaCl). Scale bars = 20 µm. All violin plots represent single-cell distributions; box plot overlays indicate medians and interquartile ranges. (H-K) Representative images and quantitative analysis of Zeiss Airyscan SR imaging for H3K27ac (H, I) and H3K4me3 (J, K) in neonatal fibroblasts following 4-hour TSA treatment at 0 nM and 250 nM. Baseline (0 nM TSA) control datasets are also presented in Fig. 2A-F and Fig. S2A-F. (H, J) Representative cross-sectional images used for quantification. White dashed boxes indicate the regions highlighted in the right composite inset panels. (I, K) Quantification of the Edge Broadening Index, defined as the average tagmentation-to-target ratio (plotted in Fig S5K, N) at the low-intensity domain edges (Target Intensity 0.1 to 0.3). For widefield imaging data (A-F), significance was determined via two-sided Mann-Whitney U tests on unclipped data for n ≥ 420 single nuclei (*p < 0.05, p < 0.01, ***p < 0.001, ****p < 0.0001, ns = not significant). Data is a representative single 96-well plate experiment of more than N ≥ 2 biological repeats. For SR imaging (I, K), significance was determined via independent Student’s t-tests on n = 5 or 6 single nuclei per condition (*p < 0.05, p < 0.01, ***p < 0.001, ns = not significant).

To isolate the spatial remodeling of the DNA from the drug-induced upregulation of the target proteins, we evaluated the tagmentation efficiency (tagmentation per unit of target). This metric explains the divergence between total intensities and localized chromatin changes. In neonatal fibroblasts, tagmentation efficiency for high-salt conditions increased at 250 nM TSA for both marks, confirming that the immediate target footprints became sterically more accessible (Fig. 4B, E). However, extending the spatial reach to 0 mM revealed distinct, target-specific dynamics. At the highest TSA dose, the H3K27ac-associated interaction hub became highly tagmentable, while the relative tagmentation efficiency for H3K4me3 plateaued or slightly decreased (Fig. 4B, E). Because H3K4me3 inherently marks focal, highly nucleosome-depleted domains at baseline, this dynamic likely reflects a local saturation effect, whereby the immediate local environment reaches maximum accessibility and yields diminishing returns for further Tn5 insertion. In contrast, the broader, more distributed domains marked by H3K27ac may experience continued physical expansion. Conversely, mature adult dermal fibroblasts exhibited a contrasting spatial response, demonstrating the assay’s sensitivity to distinct baseline compaction states. While the 300 mM footprints again became more accessible, the expanded H3K27ac local environment at 0 mM experienced a relative reduction in tagmentation efficiency, whereas the discrete H3K4me3 domains captured a progressive, dose-dependent increase in efficiency as the surrounding chromatin decondensed (Fig. 4C, F). Consistent with these metrics, direct widefield visualization confirmed that the restrictive 300 mM high-salt footprints became notably brighter following TSA treatment, while the 0 mM low-salt signals expanded into intensely fluorescent local environments (Fig. 4G).

Finally, to address the resolution limits of widefield microscopy, and to orthogonally validate that TSA induces a true physical expansion of these local environments rather than just an increase in overall nuclear fluorescence, we applied our super-resolution Airyscan pipeline to neonatal fibroblasts treated with 250 nM TSA (Fig. S5I, L). Direct visualization and globally-scaled spatial line scans confirmed increased tagmentation specifically at euchromatic domains after TSA treatment as compared to untreated baseline controls (Fig. 2C, F; Fig. 4H, J; Fig. S5I, L). Evaluating global colocalization stability (through PCC) revealed target-specific dynamics. The broadly distributed H3K27ac domains remained statistically stable across conditions (Fig. S5J), while the highly punctate H3K4me3 domains exhibited a significant increase in spatial correlation at 300 mM NaCl following TSA treatment (Fig. S5M). This further confirmed that drug-induced decondensation improves the accessibility and signal-to-noise ratio of the immediate focal footprint observed in the widefield imaging (Fig. 4E). To precisely track the spatial expansion beyond these anchors, we profiled the relative tagmentation enrichment, which revealed a dramatic spatial shift at the low-intensity edges of the protein footprints under low salt conditions (Fig. S5K, N). Quantifying this shift confirmed that TSA treatment drove a highly significant, physical expansion of the tagmentable space beyond the strict target boundary, for both H3K27ac and H3K4me3, as compared to untreated controls (Fig. 4I, K). The edge broadening index for the 300 mM high-salt condition for H3K4me3 remained statistically unchanged despite TSA treatment (Fig. 4K). This validates the biophysical rules of the assay, whereby TSA locally decondenses the chromatin to allow more frequent Tn5 insertion at the immediate anchor (reflected by the increased 300 mM widefield tagmentation efficiency), and the highly restrictive 300 mM buffer prevents the tether from physically reaching the domain margins, strictly preventing the spatial footprint from expanding.

Together, these results demonstrate that the tunable salt-stringency of tATAC-see provides a highly sensitive platform capable of isolating and quantifying dynamic spatial chromatin remodeling following pharmacological intervention.

### tATAC-see can detect chromatin remodeling at the nuclear lamina following HDAC inhibition

We next investigated whether the assay could resolve concurrent structural remodeling at the highly repressed nuclear periphery. We applied the tunable tATAC-see framework to lamin A/C and lamin B1 in both neonatal and adult dermal fibroblasts following 4-hour TSA treatment. Unlike the euchromatic targets, which exhibited simultaneous increases in both the immediate footprint and expanded local environment, the lamina targets revealed distinct, structure-specific behaviors. Upon TSA treatment, total tATAC-see intensity at 0 mM NaCl showed moderate increases, while the restrictive DNA footprint signals at 300 mM NaCl notably decreased in both cell lines (Fig. 5A, D; Fig. S6A, C, E, G). Since the target antibody intensities only mildly fluctuated (Fig. S6B, D, F, H), the reduced high-salt footprint may demonstrate the assay’s ability to detect localized loss of DNA–lamin attachment at the structural scaffold.

**Figure 5:**
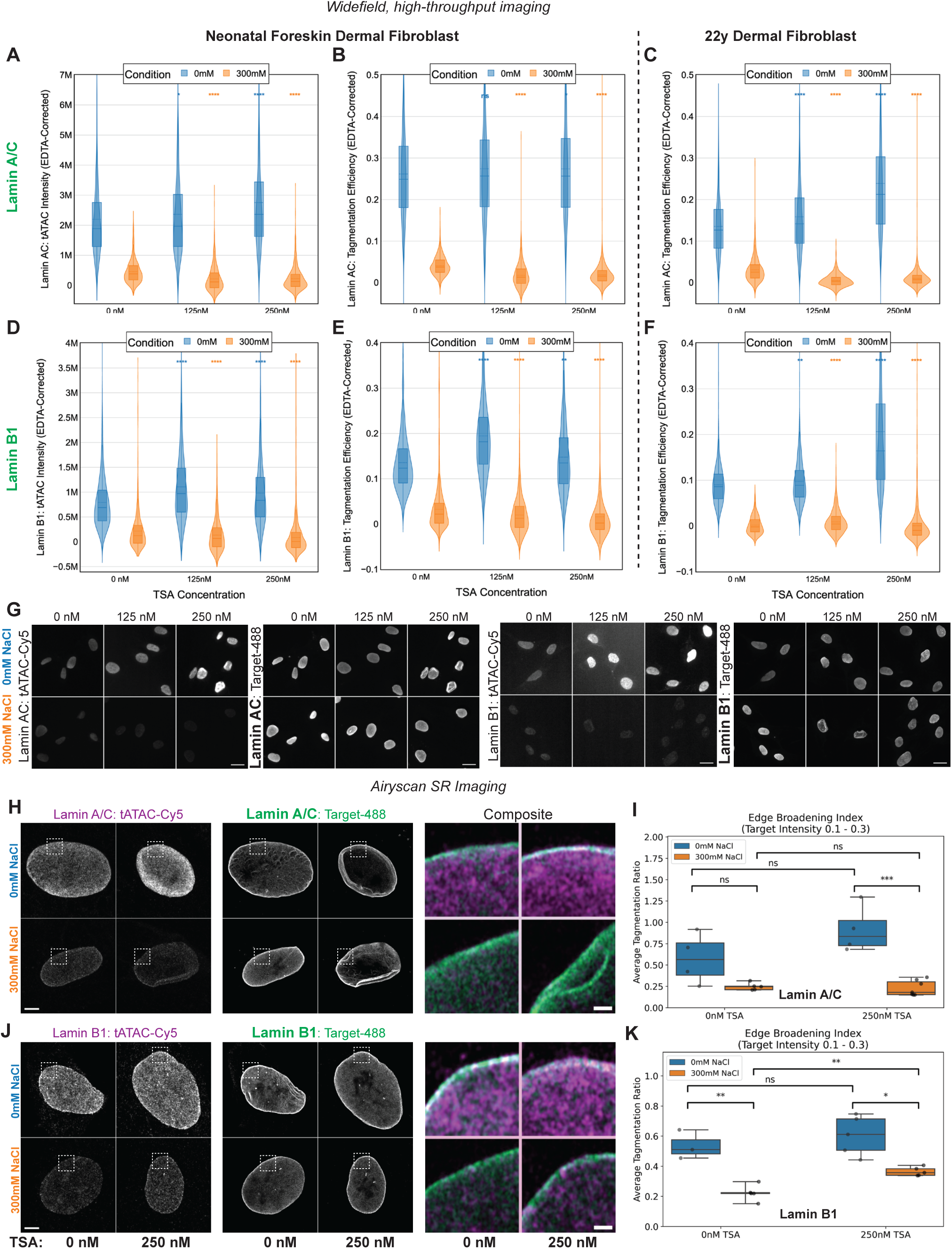
tATAC-see captures differential structural reorganization at the nuclear lamina following HDAC inhibition. (A–F) Quantitative evaluation of neonatal (A, B, D, E) and 22-year-old adult HDFs (C, F) subject to 4-hour treatment with 0, 125, or 250 nM Trichostatin A (TSA). Total tATAC-see intensity (EDTA-Corrected) (A, D) and normalized tagmentation efficiency (B, C, E, F) are plotted for lamin A/C (A-C) and lamin B1 (D-F). (G) Representative extended depth of focus (EDF) projections from widefield imaging of the tagmentation signal (tATAC-Cy5) and the structural anchor (Target-488, lamin A/C or lamin B1 primary antibody) across TSA concentrations and salt conditions (0 mM vs. 300 mM NaCl). Scale bars = 20 µm. All violin plots represent single-cell distributions; box plot overlays indicate medians and interquartile ranges. (H-K) Representative images and quantitative analysis of Zeiss Airyscan SR imaging for lamin A/C (H, I) and lamin B1 (J, K) in neonatal fibroblasts following 4-hour TSA treatment at 0 nM and 250 nM. Baseline (0 nM TSA) control datasets are also presented in Fig. 3A-F and Fig. S4A-F. (H, J) Representative cross-sectional images used for quantification. White dashed boxes indicate the regions highlighted in the right composite inset panels. (I, K) Quantification of the Edge Broadening Index, defined as the average tagmentation-to-target ratio (plotted in Fig S6K, N) at the low-intensity domain edges (Target Intensity 0.1 to 0.3). For widefield imaging data (A-F), significance was determined via two-sided Mann-Whitney U tests on unclipped data for n ≥ 420 single nuclei (*p < 0.05, **p < 0.01, ***p < 0.001, ****p < 0.0001, ns = not significant). Data is a representative single 96-well plate experiment of more than N ≥ 2 biological replicates. For SR imaging (I, K), significance was determined via independent Student’s t-tests on n = 4, 5 or 7 single nuclei per condition (*p < 0.05, p < 0.01, ***p < 0.001, ns = not significant).

To assess the chromatin reorganization at the unit level, we calculated the tagmentation efficiency. At the highest dose of TSA (250 nM), the 0 mM per-target tagmentable space increased slightly for both lamin A/C and lamin B1 in neonatal fibroblasts, but more significantly for the 22-year-old HDFs (Fig. 5B, C, E, F). The increased tagmentation efficiency indicates local decondensation at the LADs, but this may be underrepresented since a declining DNA footprint and physical detachment of chromatin from the lamina would decrease both the local Tn5 and DNA concentrations, respectively. The bell-shaped dose-response, with peak neighborhood accessibility at 125 nM TSA for lamin B1 in neonates, may support this (Fig. 5E). An initial opening of local heterochromatin may be more sensitively detected, whereas extreme hyperacetylation at higher TSA exposures may cause the DNA to loop outward, diffusing beyond the spatial reach of the tethered Tn5 anchor. Mature dermal fibroblasts show highly significant neighborhood chromatin opening and tagmentation efficiency (Fig. 5F; Fig. S6E, G), raising the possibility that alternative, cell-dependent remodeling may sterically limit DNA diffusion from the lamina. Future orthogonal structural genomic approaches (such as Hi-C or DamID) would be required to definitively confirm these localized detachment events and explore cell line–dependent differences. Divergent dose responses confirm that tATAC-see is sensitive enough to capture distinct baseline states and could be used to explore chromatin remodeling without the need for immediate sequencing.

Finally, to visually and mathematically confirm the physical expansion of LADs after HDAC inhibition, we applied our super-resolution Airyscan pipeline to neonatal fibroblasts treated with 250 nM TSA (Fig. S6I, L). Consistent with the dampened widefield efficiencies, SR imaging revealed that while the 0 mM tagmentation signal physically expanded beyond the lamina boundary upon TSA treatment (Fig. 5H, J), the magnitude of this expansion was restricted compared to euchromatic domains (Fig. 3B, E; Fig. 5H, J; Fig. S6I, L). As with control data (Fig. S4B, E), we observed that the global colocalization stability (evident through PCC) remained statistically flat, as the newly tagmented DNA did not uniformly expand around target anchors but rather likely separated towards the nuclear interior (Fig. S6J, M). Profiling the relative tagmentation enrichment again revealed muted expansions compared to euchromatin targets, supporting our widefield imaging (Fig. S6K, N). The edge broadening index therefore presented minor, if any, increases in the tagmentable space for either lamin A/C or lamin B1 as compared to untreated baseline controls (Fig. 5I, K). For lamin B1 at 300 mM, however, we observed a small but significant increase in the tagmentation ratio, opposing the expected drop observed in similar widefield metrics. This change may reflect the geometric difference between whole-nucleus widefield integration and localized optical sectioning, alongside noise from analysis under low-intensity conditions, which particularly confound SR imaging.

Overall, these data demonstrate that tATAC-see can sensitively resolve complex spatial remodeling and domain events at repressive structural anchors.

### Evaluation of tATAC-see at repressive heterochromatin targets

To define the biophysical boundaries of the tATAC-see assay, we profiled the repressive marks H3K27me3 and H3K9me3 alongside IgG negative controls using widefield imaging (Fig. S7). Direct visualization revealed that the tagmentation signal for both repressive targets prominently accumulated within the nucleoli across salt conditions (as previously observed in Fig. 1 and Fig. S1), an effect that was further exacerbated upon TSA treatment under low-salt conditions (Fig. S7A). Because TSA is known to drive the decondensation of NADs (Görisch et al., 2005), this structural relaxation likely alleviates steric hindrance, allowing the tethered Tn5 to more efficiently tagment the highly accessible nucleolar interior at 0 mM NaCl. Consequently, this spatial mislocalization confounds total nuclear image-based quantitative analysis for these specific histone marks. Nevertheless, quantitative single-cell analysis at 0 mM NaCl confirmed that the global IgG background signal was negligible after TSA treatment, verifying that the tATAC-see intensities were driven by specific antibody tethering (Fig. 4; Fig. 5; Fig. S7B-E). While H3K27me3 demonstrated significant increases in EDTA-corrected intensity and tagmentation efficiency at 250 nM TSA (Fig. S7G, I), the H3K9me3 domains appeared structurally resistant, exhibiting no significant increase (Fig. S7K, M). Notably, restrictive 300 mM high-salt signals for both heterochromatin targets were low, making robust quantitative evaluation impractical (Fig. S7B-M). This confirms, both visually and mathematically, that without the expanded 0 mM interaction radius, the steric density of the immediate heterochromatic footprint physically prevents sufficient Tn5 insertion for image-based analysis. These data establish clear technical parameters, and while tATAC-see can effectively map open neighborhoods adjacent to specific repressive anchors, applications may be fundamentally restricted by the inherent structural environment of the target.

### tATAC-see maps lamina-associated chromatin remodeling during cellular aging

Progressive chromatin opening, driven largely by the loss of peripheral heterochromatin, is a canonical hallmark of aging (López-Otín et al., 2013; O’Sullivan and Karlseder, 2012; Zhang et al., 2015). We recently demonstrated that a global chromatin opening can be captured during *in vitro* replicative and physiological aging using optimized ATAC-see (Kirkland et al., 2026). To evaluate whether tATAC-see is a practical tool for mapping complex, population-level structural shifts, we utilized these established aging models as a proof-of-principle application to quantify divergent spatial lamina-chromatin remodeling.

We first modeled *in vitro* replicative aging by culturing primary neonatal dermal fibroblasts in low, medium (mid), and high passages. We applied tATAC-see to lamin A/C, which provides structural rigidity but also interacts dynamically with chromatin (Gesson et al., 2016; Swift et al., 2013), and to lamin B1, which canonically anchors repressed constitutive heterochromatin (Steensel and Belmont, 2017). Consistent with their distinct functional roles, we observed striking, divergent biophysical trajectories. To control for cell cycle disparities between donor populations, we first tracked DNA-normalized and EDTA-corrected tATAC intensity (double-corrected) at 0 mM NaCl. Visual and quantitative analysis revealed that the total lamin A/C–associated tagmentable space progressively decreased (Fig. 6A, B), whereas total lamin B1 tagmentation moderately increased as cells were passaged (Fig. 6D, E).

**Figure 6:**
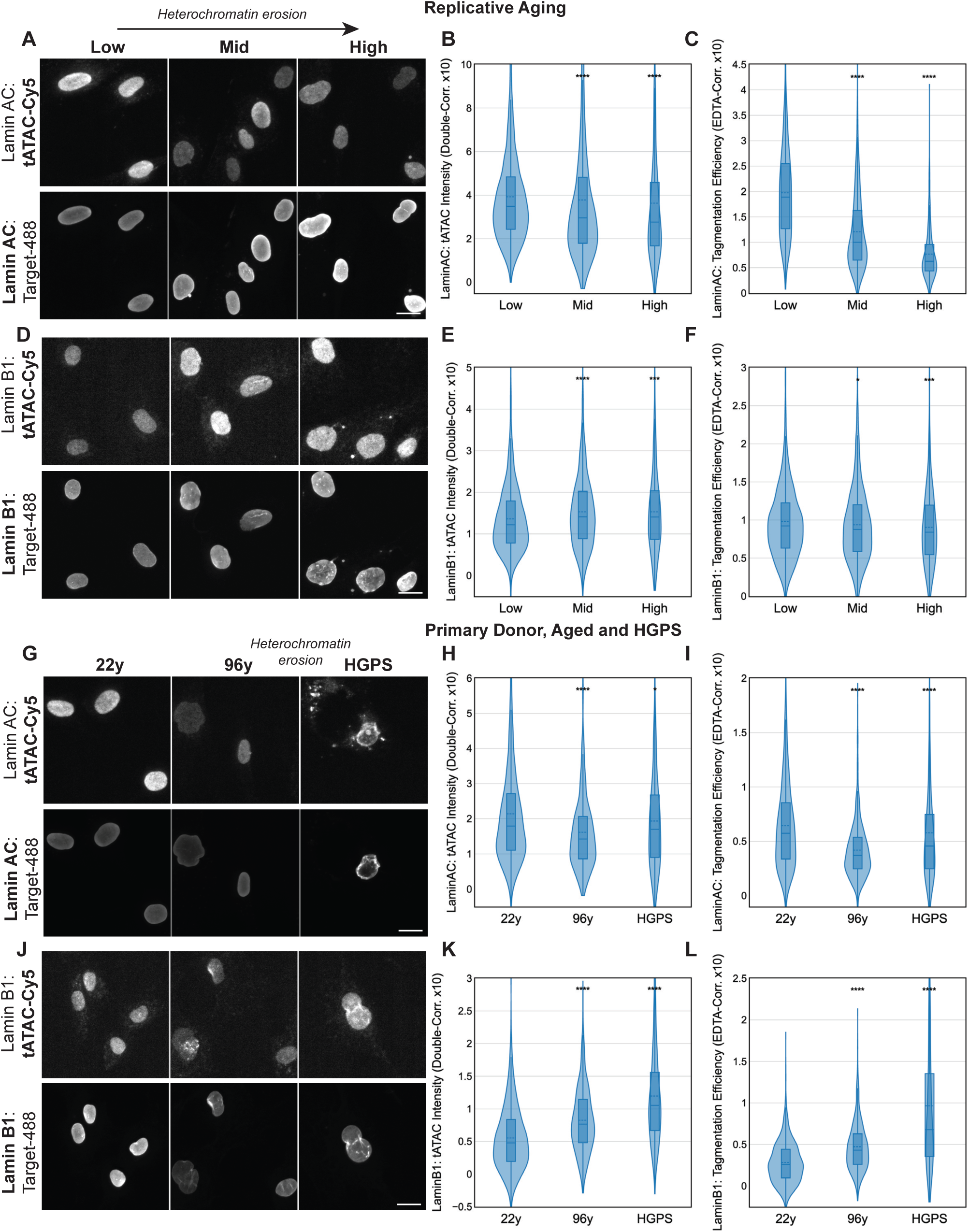
Quantitative widefield tATAC-see mapping of lamina-associated chromatin remodeling in cellular aging and progeria. (A, D, G, J) Representative extended depth of focus (EDF) projections from widefield imaging of the tagmentation signal (tATAC-Cy5) and the structural anchor (Target-488) for lamin A/C (A, G) and lamin B1 (D, J). Images display a model of replicative cellular aging across Low, Mid, and High population doublings (A, D) alongside chronological aging and Hutchinson-Gilford Progeria Syndrome models (22y, 96y, HGPS) (G, J). Scale bars = 20 µm. (B, E, H, K). Data is a representative single 96-well plate experiment of N ≥ 2 biological replicates. Total single-cell tATAC-see intensity (Double-Corrected for DNA content and EDTA background) for lamin A/C (B, H) and lamin B1 (E, K) across the corresponding aging models. (C, F, I, L) Normalized tagmentation efficiency (EDTA-Corrected) for lamin A/C (C, I) and lamin B1 (F, L) across the corresponding aging models. All violin plots represent single-cell distributions; box plot overlays indicate medians and interquartile ranges. Significance was determined via two-sided Mann-Whitney U tests on unclipped data for n ≥ 540 single nuclei per condition (*p < 0.05, p < 0.01, ***p < 0.001, ****p < 0.0001).

Notably, the decreasing lamin A/C tATAC-see intensity occurred despite a significant, passage-dependent accumulation of the lamin A/C protein anchor itself (Fig. S8C). Thus, when normalized to target abundance, tATAC-see captured a highly significant, progressive decrease in tagmentation efficiency compared to lamin B1 (Fig. 6C, F; Fig. S8A-F). Since we have validated that low-salt tagmentation conditions efficiently mark LADs, a reduction in both lamin A/C tagmentation metrics suggests chromatin is physically detaching from lamin A/C and moving out of the Tn5’s spatial radius. Importantly, this observation may capture and visually validate established genomic phenomena, where core LADs are known to actively detach and reorganize away from the nuclear envelope during cellular senescence (Lenain et al., 2017; Sadaie et al., 2013), and nucleoplasmic lamin A/C loses its interaction with euchromatic DNA as it enriches at the nuclear envelope (Scaffidi and Misteli, 2006). This physical detachment model is orthogonally supported by the decreasing lamin A/C DNA binding footprints at high-salt conditions upon passaging (Fig. S9A-C; Fig. S10A-C) and thus detects fewer lamin A/C-DNA anchor sites (Fig. 3G). On the other hand, our assay shows that lamin B1 footprints were relatively stable during *in vitro* passaging, which implies that local structural relaxation and lamin B1 protein levels may be proportionally concordant (Fig. S9D-F; Fig. S10D-F). Visually, tATAC-see intensity was maintained at sites enriched with lamin B1 staining, indicating that these specific domains may partially decondense but remained physically tethered to the periphery, whereas the results for lamin A/C appeared to show more discordance (Fig. 6A, D).

To determine if these replicative aging observations translated to human physiological and pathological aging, we applied tATAC-see to primary dermal fibroblasts derived from a 22-year-old and a 96-year-old donor, alongside fibroblasts from a patient with HGPS, a severe accelerated aging disorder driven by the toxic expression of mutant lamin A (progerin). Although analysis of a limited cohort of primary cell lines inherently carries inter-donor biological variability, using these previously characterized, passage-matched lines serves as a controlled technical validation of the assay’s capacity to measure disease- and age-associated epigenomic shifts. Concurrent with the *in vitro* aged models, the lamin A/C tagmentable neighborhood significantly decreased with chronological age, and moderately decreased in HGPS (Fig. 6G, H). Meanwhile, the lamin B1 tagmentable neighborhood dramatically increased in both the 96-year-old and HGPS cohorts (Fig. 6J, K), and this spatial relationship was conserved after accounting for the age-dependent increases in lamin A/C and reductions in lamin B1 protein levels (Fig. 6I, L; Fig. S8G-L). The high-salt conditions demonstrated that the lamin A/C DNA-binding footprints also decrease in aged-donor and HGPS fibroblasts, while lamin B1 intensities were too low to draw specific conclusions (Fig. S9G-L; Fig. S10G-L). Combined, these results propose a model reinforcing a shared trajectory of LAD remodeling in replicative, physiological and pathological aging whereby lamin A/C-associated chromatin detaches and moves beyond the reach of proximity-based tagmentation (occupying either a heterochromatic or euchromatic state), while lamin B1 domains gain relative accessibility but remain proximal to the lamin B1 anchors (Fig. 7). These findings establish the proof-of-principle that tATAC-see can identify and quantify differential biophysical chromatin shifts at specific structural anchors within the intact nucleus and uncover new biological insights for future study.

**Figure 7:**
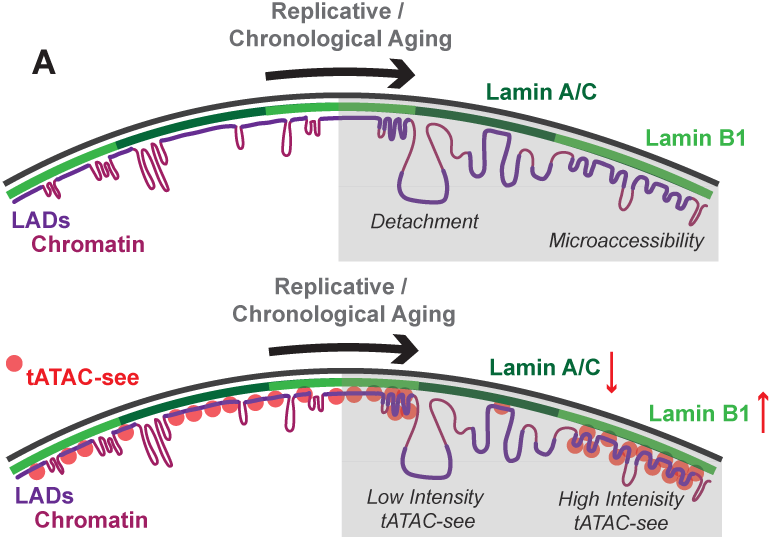
Schematic model of differential lamina-associated chromatin reorganization during aging. Proposed structural dynamics of Lamina-Associated Domains (LADs) at the nuclear periphery during replicative and chronological aging and how these may be captured with tATAC-see. In the healthy baseline state (left), compacted heterochromatin is tightly anchored to both lamin A/C and lamin B1 protein scaffolds, representing a highly constrained spatial footprint and moderate tATAC-see intensity (red spheres). Progression through cellular aging or senescence (right) drives a divergent breakdown of this peripheral architecture. Chromatin may physically detach from the lamin A/C scaffold, vacating the target-proximal local environment resulting in lowered tATAC-see intensity. Concurrently, LADs that remain anchored to the lamin B1 network may undergo structural relaxation, resulting in increased chromatin local accessibility within the localized spatial radius of the tether and elevated tATAC-see intensity.

## Discussion

In this study, we introduce targeted ATAC-see (tATAC-see), a versatile spatial epigenomic assay that merges *in situ* visualization of ATAC-see with antibody-tethered principles from CUT&Tag. By deploying this technique under low-salt conditions to expand the interaction radius of the tethered Tn5 transposase (Henikoff et al., 2020), we move beyond strict protein occupancy to visually map distinct chromatin local environments. We validated the resolution and specificity of our method using super-resolution (SR) imaging and targeted genomic sequencing, establishing a rigorous framework for interpreting rapid, accessible widefield imaging data. By determining target-dependent sensitivity to pharmacological chromatin remodeling and applying tATAC-see to models of replicative and physiological senescence, we visually captured age-dependent peripheral heterochromatin erosion. These findings establish tATAC-see as a powerful dual-modality assay that bridges spatial imaging and genomic sequencing, while providing new insight into the divergent remodeling behaviors of lamin A/C- and lamin B1-associated chromatin during aging.

### Bridging Spatial Occupancy and Chromatin Local Environments

To understand the relationship between a protein’s location and the functional accessibility of surrounding DNA, researchers typically perform independent CUT&Tag (to map protein occupancy) and ATAC-seq (to map global accessibility) assays, followed by computationally demanding multi-omic alignment (Argelaguet et al., 2021; Stuart et al., 2019). This integration is particularly challenging when relying on bulk cell populations, where rare or heterogeneous spatial variations are averaged out (Buenrostro et al., 2015; Kelsey et al., 2017). While in situ imaging techniques such as ATAC-see (Chen et al., 2016) effectively visualize the global accessible genome at single-cell resolution, they map these networks indiscriminately; they cannot identify which specific protein anchors or epigenetic marks are driving the observed topology. Currently, no spatial method can identify both where a protein is bound and resolve whether the proximal chromatin is densely compacted or highly relaxed. tATAC-see addresses this integration problem by physically profiling the local chromatin local environment surrounding a defined spatial anchor in a single *in situ* assay.

By deploying this technique under low-salt conditions (0 mM NaCl), tATAC-see functionally extends the interaction radius of the tethered Tn5 enzyme beyond the strict protein–DNA footprint. When directed against active euchromatic marks such as H3K4me3 and H3K27ac, tATAC-see visualizes the expanded spatial reach of physically proximal, interacting DNA domains (Fig. S3H). This capacity allows researchers to quantitatively track the spatial expansion of open, dynamic chromatin networks extending directly from specific regulatory anchors.

Beyond mapping established histone modifications, this capacity to profile the expanded tagmentable local environment is highly advantageous for investigating dynamic regulatory proteins. For instance, tATAC-see could be used to visualize the *in situ* activity of pioneer transcription factors, directly mapping their ability to drive structural relaxation in adjacent chromatin networks during cellular reprogramming or differentiation. Furthermore, it could be used to compare the localized structural impact of wild-type versus mutant epigenetic modifiers (such as SWI/SNF complex variants in cancer). By visually quantifying these localized chromatin remodeling events prior to sequencing, researchers can directly link specific protein binding events to the physical reorganization of local chromatin.

### Expanding ATAC Methodologies to Heterochromatic Domains

Because standard ATAC-based methods are inherently biased toward open euchromatin, they are largely ineffective for investigating densely compacted heterochromatic environments. Consequently, researchers investigating complex structural boundaries, such as LADs, typically rely on alternative methods. While optimized chromatin immunoprecipitation (ChIP-seq) (Lund et al., 2015; Sadaie et al., 2013) and tethered-nuclease assays (CUT&Run, CUT&Tag) can successfully map these insoluble heterochromatic structural anchors, they inherently require cell lysis, destroying single-cell spatial morphology, and are blind to the structural compaction state of the adjacent DNA. Furthermore, while DamID remains a historical gold standard for mapping LADs, it strictly measures the molecular contact footprint of a structural anchor; it cannot distinguish whether a tethered loop is densely compacted or locally relaxed (Steensel and Belmont, 2017). Similarly, while Hi-C sequencing can capture heterochromatin decompaction during aging, it loses single-cell spatial resolution (Criscione et al., 2016).

tATAC-see successfully bridges this methodological gap. By physically tethering Tn5 to structural anchors at the nuclear periphery, the assay generates a highly localized enzyme concentration. We observed that this targeting, particularly when combined with the expanded spatial radius permitted by the 0 mM NaCl conditions, provides a critical signal-to-noise ratio (SNR) boost that overcomes the typical steric barriers of heterochromatin. This uniquely enables ATAC-based chemistry to robustly mark LADs while retaining the spatial resolution of single-cell imaging. In our proof-of-principle application to quantify peripheral heterochromatin erosion, this allowed us to detect both the canonical detachment of LADs from lamin A/C (Lenain et al., 2017) and the localized structural relaxation of domains remaining anchored to lamin B1 during cellular aging. In addition, our data support a model in which the translocation of nucleoplasmic lamin A/C translocates to the periphery during aging (Scaffidi and Misteli, 2006), thereby depleting the dynamically associating nucleoplasmic pool and sequestering Tn5 from more accessible chromatin networks (Ikegami et al., 2020).

### Methodological Accessibility and Technical Considerations

A primary advantage of tATAC-see is its methodological accessibility. For researchers investigating spatial chromatin dynamics, the assay reduces reliance on specialized sequencing equipment and intensive bioinformatics. The protocol mirrors standard immunofluorescence (IF) workflows, enabling rapid plate-format assessment of epigenetic targets, pharmacological inhibitors, or patient-derived cell lines using standard widefield or confocal microscopy. Pairing tATAC-see with super-resolution optics allows researchers to physically trace the localized spatial topology of accessible DNA, while the preserved underlying chemistry allows for downstream genomic sequencing to identify the exact loci driving the visually observed shifts.

As with any targeted in situ method, tATAC-see possesses specific technical limitations. The assay is fundamentally dependent on highly specific, low-off-target antibodies; targets exhibiting intrinsically low signal-to-noise ratios require careful empirical optimization. Additionally, while the assay successfully profiles euchromatic networks, peripheral LADs, and dynamic targets like transcription factors, the physical bulk of the Tn5-antibody complex (∼90 kDa) experiences severe steric hindrance in the most tightly compacted constitutive heterochromatin (e.g., H3K27me3 and H3K9me3). This extreme steric exclusion can lead to off-target diffusion artifacts in domains positioned adjacent to highly accessible compartments, such as nucleoli. Because baseline nuclear permeability and compaction states vary between cell types, users must meticulously optimize the NaCl wash conditions for each new biological model to ensure that non-specifically bound Tn5 is removed prior to activation.

To maximize the utility of tATAC-see, researchers must carefully select quantitative metrics based on their specific biological questions. In this study, we distinguish between two primary readouts: total tATAC-see intensity and target-normalized tagmentation efficiency. Total intensity is the preferred metric when investigating the overall functional capacity or absolute size of a chromatin compartment (e.g., evaluating the global expansion of H3K27ac-associated regions). Alternatively, to determine whether a specific structural anchor actively remodels local chromatin, researchers should evaluate tagmentation efficiency. An increase in efficiency indicates that the binding of a transcription factor directly increases the steric availability of the local DNA per bound target molecule, offering deep mechanistic insights into local biophysics without requiring exhaustive sequencing.

With these technical considerations, tATAC-see provides a highly accessible, tunable approach for linking spatial nuclear architecture with local chromatin topology. By uncoupling the physical presence of a protein from the structural state of its local DNA, it offers a practical, powerful new framework for investigating the biophysical mechanics of aging, development, and disease.

## Methods

### Cell Culture, Passaging, and Pharmacological Treatments

Primary human neonatal dermal fibroblasts (ATCC, CRL-2522) and primary human adult dermal fibroblasts (22-year-old, 96-year-old donors, Hutchinson-Gilford Progeria Syndrome (HGPS) patient fibroblasts) (Coriell Institute, GM05294, GM00731, AG11498) were cultured in DMEM + 15% FBS + 1X NEAA (non-essential amino acids; Cat# 11140050) at 37°C in a humidified 5% CO2 incubator. All cell lines were routinely checked for mycoplasma.

For the BJ fibroblast *in vitro* aging model, cells were maintained in continuous culture and serially passaged upon reaching 70–80% confluence to prevent contact inhibition. At each passage, cells were washed with PBS and detached using TrypLE, counted using an automated cell counter, and re-seeded at a constant density of 6,600 cells/cm², equivalent to 500,000 cells per T75 flask. The aging trajectory was tracked chronologically, and representative cohorts of cells were cryopreserved at regular intervals to establish a longitudinal bank. For downstream assays, “Low passage” baseline cells were defined as those from the early expansion phase (P6–P10). “Mid passage” cells were defined as P20–P22 (approximately 4–6 weeks post-expansion), while “High passage” cells were defined as pre-senescent at P26–P28 (approximately 6 weeks of further culture). High passage cells remained proliferative, albeit at a notably reduced rate characteristic of late-stage replicative exhaustion.

For pharmacological histone deacetylase (HDAC) inhibition, cells were seeded into glass-bottom 96-well plates 24 hours prior to treatment to allow for adherence and to ensure equal cell density. Trichostatin A (TSA) (Tocris, Cat. No. 14-061) stock solutions in DMSO were diluted in culture media to final concentrations of 0, 125, or 250 nM and applied to the cells immediately following the aspiration of the standard growth media. Control cells received the equivalent dose of DMSO in media to the 250 nM treatment (0.0025%). Following a 4-hour incubation with either TSA or vehicle control, the media was removed, and cells were washed with PBS (containing CaCl_₂_ and MgCl_₂_) prior to fixation and tATAC-see processing.

Because raw single-cell fluorescence intensities cannot be perfectly normalized across independently processed imaging plates due to technical batch effects, the quantitative data and statistical distributions presented in figures are derived from a single, representative biological replicate. The trends reported were confirmed to be consistent across all independent replicates.

### Transposase Production, Assembly and Oligonucleotide Loading

#### Tn5 hyperactive double mutant with n-terminal protein-a/g and c-terminal mxe intein and chitin binding domain

MSLKDDPSQSANLLSEAKKLNESQAPKADNKFNKEQQNAFYEILHLPNLNEEQRNGFIQSLKDDPSQSANLLAEAKKLNDAQAPKADNKFNKEQQNAFYEILHLPNLTEEQRNGFIQSLKDDPSVSKEILAEAKKLNDAQAPKTTYKLVINGKTLKGETTTEAVDAETAERHFKQYANDNGVDGEWTYDDATKTFTVTEKPEVIDASELTPAVDDDKEFGGGGSGGGGSGGGGSGGGGSHMITSALHRAA DWAKSVFSSAALGDPRRTARLVNVAAQLAKYSGKSITISSEGSKAMQEGAYRFIRNPNVSAEAIRKAGAMQTVKLAQEFPELLAIEDTTSLSYRHQVAEELGKLGSIQDKSRGWWVHSVLLLEATTFRTVGLLHQEWWMRPDDPADADEKESGKWLAAAATSRLRMGSMMSNVIAVCDREADIHAYLQDKLAHNERFVVRSKHPRKDVESGLYLYDHLKNQPELGGYQISIPQKGVVDKRGKRKNRPARKASLSLRSGRITLKQGNITLNAVLAEEINPPKGETPLKWLLLTSEPVESLAQALRVIDIYTHRWRIEEFHKAWKTGAGAERQRMEEPDNLERMVSILSFVAVRLLQLRESFTPPQALRAQGLLKEAEHVESQSAETVLTPDECQLLGYLDKGKRKRKEKAGSLQWAYMAIARLGGFMDSKRTGIASWGALWEGWEALQSKLDGFLAAKDLMAQGIKICITGDALVALPEGESVRIADIVPGARPNSDNAIDLKVLDRHGNPVLADRLFHSGEHPVYTVRTVEGLRVTGTANHPLLCLVDVAGVPTLLWKLIDEIKPGDYAVIQRSAFSVDCAGFARGKPEFAPTTYTVGVPGLVRFLEAHHRDPDAQAIADELTDGRFYYAKVASVTDAGVQPVYSLRVDTADHAFITNGFVSHATGLTGLNSGLTTNPGVSAWQVNTAYTAGQLVTYNGKTYKCLQPHTSLAGWEPSNVPALWQLQ

#### pAG-Tn5 Expression

The gene encoding the Protein A/G-tagged Tn5 double mutant (pAG-Tn5) was synthesized and cloned into empty pET3a vectors (Sigma-Aldrich, Cat. No. 69418) using NdeI and BamHI restriction sites. The plasmid was transformed following standard protocols into BL21(DE3) *E. coli* (Thermo Fisher Scientific, Cat. No. EC0114) and grown at 37°C overnight on LB agar plates containing 100 µg/mL ampicillin (Teknova, Cat. No. L1002). An 80 mL LB starter culture containing 100 µg/mL carbenicillin (Corning, Cat. No. 46-100-RG) in a 250 mL baffled flask was inoculated with a single isolated colony and grown at 30°C overnight. The following morning, 10 mL of dense starter culture was sub-cultured into 1 L of LB containing 100 µg/mL carbenicillin in a 2.7 L baffled flask. Cultures were grown at 37°C until reaching an optical density (OD_₆₀₀_) between 0.6 and 0.8. Protein induction was subsequently initiated by the addition of 1 mM IPTG (Thermo Scientific, Cat. No. 34060), and the incubator temperature was reduced to 22°C for 18 h. Cells were harvested by centrifugation at 8,000 x *g* using a JLA-8.1000 rotor in an Avanti centrifuge (Beckman Coulter), and cell pellets were stored at −80°C until purification.

#### Tn5 Lysis and Purification

For lysis, cell pellets were thawed on ice for approximately 2 h and resuspended in 10 mL per gram of HEGX buffer (50 mM HEPES pH 7.2, 800 mM NaCl, 0.8 mM EDTA, 10% v/v glycerol, 0.2% Triton X-100) supplemented with protease inhibitor cocktail (Sigma-Aldrich, Cat. No. P8849). Following complete resuspension, cells were disrupted by sonication on ice using a 5 sec on / 5 sec off cycle at 60% amplitude for 10 min total sonication time. The lysate was transferred to centrifuge tubes and cleared at 40,000 x *g* for 20 min at 4°C using a JA-25.50 rotor. The supernatant was decanted into a beaker, and bulk DNA was precipitated by the dropwise addition of 10% v/v polyethylenimine (PEI) (Sigma-Aldrich, Cat. No. 482595) to a final concentration of 0.1% v/v, or until the solution became visibly cloudy. The solution was centrifuged again at 40,000 x *g* for 20 min at 4°C. The clarified supernatant was decanted from the DNA pellet for immediate purification.

The subsequent purification steps were performed entirely at 4°C using pre-chilled buffers. The clarified supernatant was applied to a gravity-flow column containing 0.5 mL of HEGX-equilibrated chitin resin (New England Biolabs, Cat. No. S6651L) per gram of lysed cell pellet. The lysate was passed through the column twice to ensure maximal binding. The resin was washed with 30 column volumes (CV) of HEGX buffer. Intein cleavage was initiated by the addition of 2 CV of elution buffer (HEGX containing 100 mM freshly dissolved DTT). One CV of elution buffer was allowed to flow through the column before the stopcock was closed; the remaining 1 CV of buffer was incubated on the column for 72 h to ensure complete intein cleavage and Tn5 liberation. Following the 72 h incubation, the elution buffer was collected, and the column was washed with an additional 1 CV of HEGX to recover residual liberated Tn5.

The eluted protein was concentrated to 10 mL and injected onto a HEGX-equilibrated 320 mL HiLoad 16/600 Superdex 200 pg size exclusion column (Cytiva, Cat. No. 28989335) utilizing an ÄKTA pure FPLC system (Cytiva). The column was washed with 1 CV of HEGX, and fractions were collected via an automated fractionator. The Tn5 peak eluted at approximately 0.6–0.8 CV, and all corresponding fractions were pooled. Total protein concentration was determined via BCA assay (Thermo Scientific, Cat. No. 23225) and concentrated to a final target of 2.5 mg/mL (approximately 27 µM). The protein was dialyzed overnight into storage buffer (50 mM HEPES, 800 mM NaCl, 0.2 mM EDTA, 20% v/v glycerol, 2 mM DTT), aliquoted into microcentrifuge tubes (256 µL per tube), flash-frozen in liquid nitrogen, and stored at −80°C.

### Mosaic End Sequences and Transposome Loading

Single-stranded DNA oligonucleotides were synthesized by Integrated DNA Technologies (IDT) and dissolved to stock concentrations of 200 µM using nuclease-free water. The sequences utilized were as follows, where the 5’ Cy5 fluorophore can be substituted with Cy3 or Alexa Fluor 488 depending on experimental requirements:

- Tn5ME-Reverse (R): 5’-[Phos] CTG TCT CTT ATA CAC ATC T -3’
- Tn5ME-A-Cy5 (A): 5’-[Cy5] TCG TCG GCA GCG TCA GAT GTG TAT AAG AGA CAG -3’
- Tn5ME-B-Cy5 (B): 5’-[Cy5] GTC TCG TGG GCT CGG AGA TGT GTA TAA GAG ACA G -3’

To prepare 1,028 µL of 20X (3.4 µM) fluorescently loaded transposomes for tATAC-see, double-stranded adapter complexes were first generated. The A-R complex was created by combining 12.8 µL of 200 µM Tn5ME-A-Cy5 with 12.8 µL of 200 µM Tn5ME-Reverse in 25.6 µL of annealing buffer (10 mM Tris-HCl, pH 8.0) to achieve a 50 µM total DNA concentration. The B-R complex was similarly prepared by combining 12.8 µL of 200 µM Tn5ME-B-Cy5 with 12.8 µL of 200 µM Tn5ME-Reverse in 25.6 µL of annealing buffer. The A-R and B-R oligonucleotides were annealed in separate PCR tubes using a T100 thermal cycler (Bio-Rad) programmed to heat to 95°C for 5 min, cool to 65°C at a rate of 0.1°C/sec, hold at 65°C for 5 min, and finally cool to 4°C at 0.1°C/sec.

The annealed A-R and B-R oligonucleotides were independently diluted to 20 µM by adding 76.8 µL of an 85% v/v aqueous glycerol solution to each 51.2 µL reaction. Both solutions (128 µL each) were combined into a single 1.5 mL microcentrifuge tube, yielding 256 µL of mixed adapters. This mixture was further diluted with 640 µL of 50% v/v glycerol, bringing the volume to 886 µL. Finally, 128 µL of the thawed 27 µM pAG-Tn5 stock was added. The solution was mixed thoroughly by pipetting and incubated for 1 h at room temperature in the dark to allow for complete transposome loading. The final fully loaded transposome complex (1,028 µL in 47% v/v glycerol) was stored at −30°C until experimental use.

### Targeted ATAC-see (tATAC-see) Assay

For imaging assays only, cells were seeded at a density of 5000 cells per well into 96-well glass-bottom plates, for combined imaging and sequencing assays, 100,000 cells were seeded per well of 12-well glass-bottom plates and cultured for 24 hours at 37°C and 5% CO_₂_ under standard growth conditions to allow for firm cellular attachment and morphological recovery. To begin the assay, cells were rinsed with Hank’s Balanced Salt Solution (HBSS) containing calcium and magnesium (Fisher Scientific, Cat. No. 14065056) and lightly fixed using 1% paraformaldehyde (PFA) (Electron Microscopy Sciences, Cat. No. 15714) in HBSS containing calcium and magnesium for 10 minutes at room temperature. Following fixation, cells were washed three times with HBSS containing calcium and magnesium. Plates were now stored at 4C, or used immediately for tATAC-see. HBSS <u>without</u> calcium and magnesium (Fisher Scientific, Gibco, Cat. No. 4185052) was now used from this point. First, cells were permeabilized using 0.5% Igepal (Abcam, Cat. No. ab285400) in HBSS for 15 minutes at room temperature. To minimize non-specific binding, cells were blocked for 60 minutes at room temperature in a blocking buffer consisting of 2% BSA (Sigma-Alrich, Cat. No. A7906) and 0.1% Igepal in HBSS. Cells were subsequently incubated with primary antibody at appropriate concentrations prepared in blocking buffer for 1-2 hours at room temperature or overnight at 4C. Cells were washed with HBSS containing 0.1% Igepal, three times for 10 minutes. Next, cells were incubated with appropriate secondary antibodies, diluted in blocking buffer for 60 minutes at room temperature. Prior to enzyme tethering, cells were again washed three times for 10 minutes, each in 0.1% Igepal in HBSS. For spatial targeting, cells were incubated with Cy5-loaded pAG-Tn5 (1:100 dilution, 34nM) in a tethering buffer containing 1% BSA, 0.1% Igepal, 300 mM NaCl (total with HBSS, thus 163 mM added) (Teknova, Cat. No. S0250), 2.5 mM EDTA (Invitrogen, via Fisher Scientific, Cat. No. AM9262), and HBSS for 1 hour at room temperature. Unbound enzyme was strictly removed via three 10-minute high-salt washes consisting of 300 mM NaCl, 2.5 mM EDTA and 0.1% Igepal in HBSS. Note: When establishing new targets and cell lines, perform a visual check to ensure removal of non-specific DNA bound Tn5 for IgG controls, and specific signal for targets. Increased salt concentration or longer washes may be necessary. A quick rinse was performed using 0.1% Igepal in HBSS with 300 mM NaCl to dilute residual EDTA before adding TD buffer prepared from a 2X stock with final TD concentrations: 10 mM TAPS, pH 8.0, 5 mM MgCl_₂_, 10% *N,N*-dimethylformamide, 1% BSA. EDTA was added at 20 mM for background control samples, and appropriate volumes of NaCl for salt titration and DNA footprint experiments. Plates were then incubated at 37°C for 1 hour with optional gentle agitation. All tagmentation reactions were immediately quenched for 5 minutes using a stop buffer containing 300 mM NaCl, 0.01% SDS (Invitrogen, via Fisher Scientific, Cat. No. 15-553-027), and 40 mM EDTA. Cells were subsequently rinsed with HBSS three times before nuclei were counterstained with 1:1000 Hoechst 33342 (Invitrogen, via Fisher Scientific, Cat. No. H3570) for 25-minutes to enable nuclear segmentation, and rinsed again three times with HBSS.

#### Antibodies for in situ Spatial Targeting

The primary antibodies and dilutions utilized were: rabbit anti-lamin A/C (Proteintech, Cat. No. 10298, 1:400), rabbit anti-lamin B1 (Proteintech, Cat. No. 12987, 1:500), rabbit anti-H3K27ac (Abcam, Cat. No. ab4729, 1:1000), anti-H3K27me3 (Millipore, Cat. No. 07-449, 1:200), mouse anti-H3K4me3 (Active Motif, Cat. No. 61379, 1:500), mouse anti-H3K9me3 (Active Motif, Cat. No. 61013, 1:400), and an IgG control (Invitrogen via Fisher Scientific, Cat. No. PI31235, 1:500). Following primary incubation and washing, cells were incubated with secondary antibodies: mouse anti-rabbit FITC (Invitrogen, Cat. No. 31584, 1:1000) and rabbit anti-mouse 488 (Invitrogen, Cat. No. A-11059], 1:2000) in 2% BSA blocking buffer for 1 hour at room temperature. Mouse and Rabbit secondaries boosted the binding of pA/G-Tn5.

#### Tunable Salt-Gradient Tagmentation Optimization

To systematically evaluate the spatial radius of the tethered tagmentation across varying chromatin domains, neonatal foreskin dermal fibroblasts (BJ) and 22-year old HDFs (GM05294) were subjected to a tagmentation salt gradient. During the tagmentation step of the general protocol, the reaction buffer was formulated with the following concentrations of NaCl: 300 mM, 150 mM, 75 mM, and 0 mM. To establish a strict baseline for non-specific background, negative control wells for both the 300 mM and 0 mM NaCl conditions were supplemented with 20 mM EDTA to chelate essential ions and completely inhibit enzymatic activity.

### Global ATAC-see

Global ATAC-see was performed as described in (Kirkland et al., 2026) with only plate format alterations for complimentary sequencing assays. Briefly, 100,000 cells were cultured in a 12-well glass bottom plate (Cell Vis) 24 hours prior to processing. Cells were washed once with HBSS containing calcium and magnesium (HBSS + ions; Thermo Fisher, Cat# 14025134) and fixed using 1 ml of freshly diluted 1% (w/v) paraformaldehyde (PFA; Cat# 50-980-494) in HBSS + ions for 10 min at room temperature. Wells were washed with 2 ml of HBSS without calcium and magnesium (HBSS w/o Ions; Cat# 14065056), three times. Plates were immediately processed or stored at 4°C for up to one week. Transposomes (3.4 µM stock) were diluted 20-fold to a final working concentration of 170 nM in 1X Tagment DNA (TD) buffer (10 mM TAPS, pH 8.0, 5 mM MgCl_₂_, 10% *N,N*-dimethylformamide; prepared from a 2× stock) supplemented with 0.1% IGEPAL CA-630, and mixed thoroughly by pipetting. The working transposome solution was sonicated in a bath sonicator at 37 kHz for 60 seconds at 80% amplitude and passed through a 0.22 µm syringe filter (Cat# SLGVR33RS) to remove residual aggregates. The tagmentation solution was applied to aspirated wells (1 ml for a single well in a 12-well plate) and plates were incubated for 60 minutes at 37°C. Wells were aspirated and the tagmentation reaction was quenched by washing the cells with 2 ml of HBSS w/o ions, supplemented with 40 mM EDTA, 163 mM NaCl (final concentration equal to 300 mM NaCl) and 0.01% SDS. Wells were rinsed three times with HBSS w/o ions and stained with Hoechst for 25-minutes to enable nuclear segmentation and rinsed again three times with HBSS w/o ions. Plates were stored at 4°C or imaged immediately and subsequently processed for sequencing.

### High-Throughput Widefield Imaging

Automated widefield epifluorescence imaging was performed utilizing a Nikon Eclipse Ti2 inverted microscope integrated into a BioPipeline high-throughput imaging system. Prepared 96-well plates were automatically loaded and scanned using a Nikon CFI Plan Apochromat Lambda D 20X air objective (NA 0.80) paired with a Photometrix Kinetix 25mm camera, Lumencor Spectra III light engine, Semrock Brightline individual or seven-filter “Pinkel” penta-band filter cubes, and a Cairn OptoSpin filter wheel with individual 32mm Semrock Brightline emission filters. To ensure a robust dynamic range for the quantitative assessment of both intense localized foci and diffuse nucleoplasmic signals, all images were captured at a 16-bit pixel depth. Multi-channel acquisition was fully automated via NIS-Elements software. To compensate for micro-fluctuations in plate topography during automated scanning, Nikon’s Perfect Focus System (PFS) was employed to establish a stable baseline focal plane. Images were acquired at multiple spatially distributed fields of view per well. At each field of view, images were captured as Z-stacks consisting of 9 optical slices with a 0.8 µm step size.

### Airyscan SR Imaging

Super-resolution (SR) imaging was performed using a Zeiss LSM 980 confocal microscope equipped with an Airyscan detector. Images were acquired using a Plan-Apochromat 63x/1.4 NA Oil DIC M27 objective with a 4x optical zoom. To satisfy Nyquist sampling criteria, optimal frame sizes (768 x 768 pixels) were utilized, yielding a voxel resolution of 0.043 × 0.043 × 0.17 µm. Images were captured at a 16-bit depth to ensure a high dynamic range for quantitative intensity measurements. Acquisition was performed utilizing unidirectional scanning at a scan speed of 7 (3.89 s frame time), and 2x line averaging was applied to optimize the signal-to-noise ratio. Prior to the acquisition of each field of view, the Airyscan detector alignment was optimized. Raw multiplexed images were subjected to batch 3D Airyscan processing and automated channel alignment using ZEN Blue software (Carl Zeiss) to resolve sub-diffraction structures and correct for three-dimensional chromatic aberrations.

For quantitative spatial analysis, automated image processing pipelines were utilized to systematically extract the optical cross-section of each nucleus. To eliminate manual selection bias, a custom script was applied to evaluate the entire Z-stack and automatically extract the single Z-slice exhibiting the maximum mean target fluorescence, representing the focal center of the nucleus.

### Image Processing and Quantitative Analysis

#### Processing and quantification of widefield epifluorescence microscopy

For automated widefield epifluorescence imaging was performed utilizing a Nikon Eclipse Ti2 inverted microscope, to establish a standardized 2D focal plane from the acquired multi-slice Z-stacks, raw images were processed into single-plane projections utilizing the Extended Depth of Focus (EDF) algorithm within Nikon NIS-Elements software. Subsequent image analysis and single-cell feature extraction were performed using Arivis Pro 4.2.0 and 4.3.0 software. Single-nucleus segmentation was achieved by applying the integrated Cellpose deep-learning tool (utilizing the pre-trained cyto3 model) directly to the Hoechst counterstain channel. The verified nuclear masks were then applied across all corresponding experimental channels to extract single-cell morphological and fluorescence intensity metrics. Exported CSV files were processed in Python 3.11.4 using Plotly, Seaborn, Pandas and Numpy libraries. Statistical tests were executed using the Scipy.Stats module.

To establish the baseline of non-specific oligonucleotide retention, matched EDTA-inhibited control samples were processed for each target and salt condition (0 mM and 300 mM NaCl). For establishing the specificity of the assay, the median background signal from the EDTA controls was calculated per target and subtracted from the corresponding active enzyme samples. To preserve the natural distribution of technical noise and prevent artificial deflation of sample variance, background-subtracted values were left unclipped (allowing negative noise values) for all statistical testing. For logarithmic data visualization only, values were clipped at a minimum of zero to prevent mathematical distortion of the lower bounds. For the visualization of background-subtracted ATAC intensities spanning multiple orders of magnitude, a symmetrical log (SymLog) transformation was applied. This shifted logarithmic scale compresses the optical noise floor linearly while preserving the logarithmic separation of high-intensity biological signals, preventing visual distortion of the baseline distribution. Due to an aggregation artifact in the specific EDTA-inhibited control well for H3K27ac in Neonatal Fibroblasts treated with 250 nM TSA, the median background signal from the adjacent 125 nM TSA EDTA-inhibited H3K27ac condition was utilized as a proxy baseline. To compare independent cell lines, raw tATAC intensities from both experimental and EDTA control samples were normalized by the respective DNA intensity per cell before and median EDTA subtraction to control for cell cycle and background intensity respectively.

To account for target abundance and assess DNA tagmentation per unit of target, "Tagmentation Efficiency" was calculated by dividing the ATAC signal by the target antibody signal for each individual cell. To combine background correction from non-specific signal or retention of Tn5 with bound fluorescent oligonucleotides, we subtracted the tagmentation efficiency of the EDTA ATAC from the raw ATAC ((raw ATAC / Target) - median of EDTA ATAC / Target)).

Due to the non-normal distribution typical of single-cell intensity data, all statistical comparisons between conditions were performed using a two-sided Mann-Whitney U test. Significance thresholds were defined as: \**p* < 0.05*, **p < 0.01, ***p < 0.001*, and \*\*\*\**p* < 0.0001. Because raw tagmentation efficiency ratios cannot be directly compared across different targets due to variable antibody binding affinities and fluorophore conjugation efficiencies, cross-target comparisons of chromatin expansion were evaluated using Cohen’s *d* effect size. Cohen’s *d* was calculated using the pooled standard deviation to provide a standardized, dimensionless measure of the magnitude of accessibility gain between the 0 mM and 300 mM states, completely independent of absolute antibody brightness. A Cohen’s *d* effect size > 0.2 denoted expansion, and d > 0.8 is conventionally considered a large effect.

To rigorously validate tagmentation expansion independent of metric scale or ratio-based artifacts (utilized for data visualization in Supplementary Fig. S1), data were transformed into Modified Z-scores. Unlike standard Z-scores which rely on the mean and standard deviation, the Modified Z-score utilizes the median and the Median Absolute Deviation (MAD), rendering it highly resistant to single-cell outliers. The Modified Z-score (*M_i_*) for each cell was calculated using the formula:

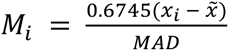

where *x*∼ is the median of the 300 mM baseline condition, and MAD is the median absolute deviation of the 300 mM baseline. To ensure the true technical noise floor was captured, the MAD was calculated using unclipped background-subtracted data. A minimal floor was applied to the MAD to prevent undefined values in cases of extreme target drop-off. Consequently, the 300 mM condition centers precisely at *M_i_* = 0, expressing the 0 mM signal as the robust standard deviations of expansion above the technical baseline.

To account for cell cycle variability and global senescence-associated spatial dilution, DNA-normalized tATAC-see intensity was calculated by dividing the raw tATAC-see signal by the cell matched Hoechst intensity.

#### Processing and quantification of SR airyscan microscopy

For quantitative spatial analysis of airyscan SR images, automated image processing pipelines were first utilized to systematically extract the optical cross-section of each nucleus. To eliminate manual selection bias, a custom script was applied to evaluate the entire Z-stack and automatically extract the single Z-slice exhibiting the maximum mean target fluorescence, representing the focal center of the nucleus.

Dual-channel microscopy images were then processed using a custom automated Python pipeline (scikit-image 0.26.0, scipy 1.17.1). To restrict the analysis strictly to nuclear regions, a binary exclusion mask was generated for each image by applying a Gaussian blur (σ = 2) to the target chromatin channel, followed by Li’s minimum cross-entropy thresholding. Objects smaller than 500 pixels were discarded, and internal holes were filled to create a uniform, solid nuclear mask. To enable robust quantitative comparisons across independent experimental conditions (e.g., varying salt and TSA concentrations), a dual-global normalization strategy was employed. The absolute 1st and 99th intensity percentiles within the nuclear masks were calculated across the entire combined dataset. Every image channel was subsequently min-max normalized to these universal global minima and maxima. This locked all fluorescence values to a fixed scale from 0 to 1, preserving the relative quantitative loss or gain of spatial signal across all biological conditions.

Global colocalization was assessed by calculating the Pearson Correlation Coefficient (PCC) of the normalized pixel intensities within the nuclear mask. For spatial enrichment profiling, the normalized target pixel intensities were sorted into 20 equal bins (0.0 to 1.0). For each bin, the relative tagmentation enrichment was calculated as the ratio of the mean tagmentation signal to the median target bin intensity. Cell profiles were averaged, and the Standard Error of the Mean (SEM) was calculated to visualize population variance. To quantify spatial expansion, an "Edge Broadening Index" was defined as the average tagmentation ratio within the low-intensity target chromatin bins (0.1 to 0.3 normalized intensity). The extreme nuclear periphery (0.0 to 0.1) was deliberately excluded to prevent mathematical inflation of the ratio caused by near-zero target values and background noise. Statistical significance across the more than two experimental conditions was first evaluated using a global one-way ANOVA. For specific condition comparisons, independent Student’s t-tests were performed. Multiple tests were subjected to a Bonferroni correction to account for multiple hypothesis testing (*n* = 4 planned comparisons). Adjusted p-values are reported as: *p < 0.05, **p < 0.01, ***p < 0.001, and ns = not significant.

As single-cell segmentation and intensity extraction for both widefield and SR imaging were executed using these fully automated computational pipelines, user blinding during image acquisition and analysis was not required. Statistical power analyses were not performed *a priori* to determine sample sizes.

### Sequencing Library Preparation and Data Analysis

Following imaging on a Nikon Eclipse Ti2 inverted microscope as described above, reverse cross-linking was performed as in and performed as in (Kirkland et al., 2026). Briefly, HBSS was aspirated and replaced with a reverse-crosslinking solution adapted from (Chen et al., 2016) consisting of 50 mM Tris-HCl, 1 mM EDTA, 1% SDS, 0.2 M NaCl and 250 ug/ml proteinase K. Cells were physically detached using a cell scraper, transferred to microcentrifuge tubes, and incubated overnight at 65°C with agitation at 1,000 rpm in a thermomixer. DNA was subsequently purified using the MinElute Reaction Cleanup Kit (Qiagen, Cat#: 28204) according to the manufacturer’s instructions. Eluted DNA was amplified using 11, 14, and 17 PCR cycles with NEBNext® HiFi 2× PCR Master Mix (NEB, M0541S) and pre-mixed Illumina DNA/RNA UD Indexes (Set A; cat. no. 20091646), following the PCR protocol described in (Buenrostro et al., 2013). Library quality was assessed by Agilent TapeStation, and products from all three cycle numbers were pooled and purified using 1.8X SPRI beads (Beckman Coulter, cat. no. B23318). Sequencing was performed on an Illumina NextSeq 2000 (P3 flow cell) with paired-end 50 + 50 bp reads and 10 bp dual-indexing (50/10/10/50 cycle configuration). Reads were processed with nf-core/atacseq (Ewels et al., 2020) (v2.1.2), aligned to GRCh38 and peaks called with MACS2 (Zhang et al., 2008), yielding narrowPeak files.

Peaks were annotated with ChIPseeker (Yu et al., 2015) against the UCSC hg38 known-gene model (TxDb.Hsapiens.UCSC.hg38.knownGene; gene symbols from org.Hs.eg.db); feature proportions are shown as pie charts.

lamin A/C and lamin B1 Targeted ATAC-see coverage at 300 mM and 0 mM was expressed as a log2 ratio over the BJ IgG 0 mM control with deepTools (Ramírez et al., 2016) bamCompare (10-kb bins, 30-kb smoothing, read-count scaling, reads extended to fragment length, duplicates ignored, GRCh38 effective genome size 2.91 × 10⁹ bp).

For each mark, peaks from the 0 mM, 300 mM and Global 0 mM sets were merged per chromosome into non-overlapping consensus regions (overlapping or directly abutting intervals joined), and each region was assigned to the combination of sets contributing at least one peak, giving the seven membership classes of the UpSet (Lex et al., 2014) plot (drawn in Python).

Distances were computed between peak summits (narrowPeak point-source offset, or interval midpoint where absent) within chromosomes. Gained peaks were defined as 0 mM peaks overlapping no same-mark 300 mM peak (21,524 for H3K4me3, 60.2%; 40,268 for H3K27ac, 39.4%). For each gained peak, the distance to the nearest 300 mM summit was compared with that of a non-anchor accessibility background (Global 0 mM peaks not overlapping a 300 mM peak; n = 127,659 for H3K4me3, 79,672 for H3K27ac) and a uniform null (gained summits randomized within each chromosome’s peak span).

Distance distributions were compared by one-sided Mann–Whitney U test with Cliff’s δ as the effect size (H3K4me3 δ = +0.21; H3K27ac δ = +0.16; both P ≈ 0). Given the large sample sizes (10^⁴^–10^⁵^), effect sizes and the distance-resolved enrichment are emphasized over nominal P values. A fixed random seed was used for all randomizations and bootstrapping.

Overlap, distance and locality analyses and all custom figures were implemented in Python 3.12 with NumPy (Harris et al., 2020), pandas (Virtanen et al., 2020), SciPy (McKinney, 2010) (statistical tests) and Matplotlib (Hunter, 2007); coverage, log2-ratio and metagene analyses used deepTools (Lex et al., 2014); peaks were called with MACS2 (Zhang et al., 2008) within nf-core/atacseq (Ewels et al., 2020); annotation used ChIPseeker (Yu et al., 2015); tracks were viewed in IGV (Robinson et al., 2011).

## Use of Artificial Intelligence tools

The large language model Gemini (Google) was used to adapt code for data visualization and to refine manuscript text. The AI was not used for data analysis or scientific interpretation. The authors have reviewed and verified all AI-assisted code, text, and references for accuracy and take full responsibility for the content of this manuscript.

## Acknowledgements

We would like to thank Eric Griffis and Judy Martin of the Imaging hub, and Tatyana Makushok, Maria Burdyniuk, Valentina Pedoia from the Biovision Hub for supporting microscopy and image analysis training. We also thank Christa Caneda and Garret Bishop of the Cell Modeling and Engineering Hub for supporting all *in vitro* operations and we thank Nasun Ha and the genomics hub for supporting sequencing. Finally, we thank the operations and facilities teams for enabling all daily activities. We thank Francesco Della Valle for comments on the manuscript.

## Author Contributions

N.K., Y.Y. and M.A.C. conceived the project, performed the core experiments, and wrote the original manuscript. N.K. and Y.Y. analyzed and interpreted the quantitative data. M.A.C. designed and conducted the biochemical protocols for Tn5 expression and purification. P.M-C. critically reviewed and edited the manuscript, Z.L. supervised the research program.

## Competing Interests

All authors are current employees of Altos Labs and hold equity (shares and/or options) in the company.

## Funding

This study was funded by Altos Labs, Inc. All authors are employees of Altos Labs, Inc.

## Data and Resource availability

The raw and processed sequencing data generated and analyzed in this study have been deposited in the NCBI Gene Expression Omnibus (GEO) under accession number GSE338034 and GSE338065. Due to proprietary institutional restrictions, raw imaging datasets are not publicly deposited but are available from Altos Labs, Inc. upon reasonable request for non-commercial research purposes, subject to a standard institutional Data Sharing Agreement (DSA).

Custom image analysis workflows were executed using commercially available software (Nikon NIS-Elements, Arivis Vision4D) as described in the Methods. Custom Python scripts and Jupyter Notebooks utilized for spatial metrics, statistical analysis, and data visualization will be made available upon reasonable request for non-commercial research purposes, subject to a standard institutional Data Sharing Agreement (DSA).

## Notes

### Summary of Updates

Update author information and pdf conversion issues

