## Supplemental Figures for "Targeted ATAC-see (tATAC-see): A Visual Assay for Target-Specific Chromatin Profiling"

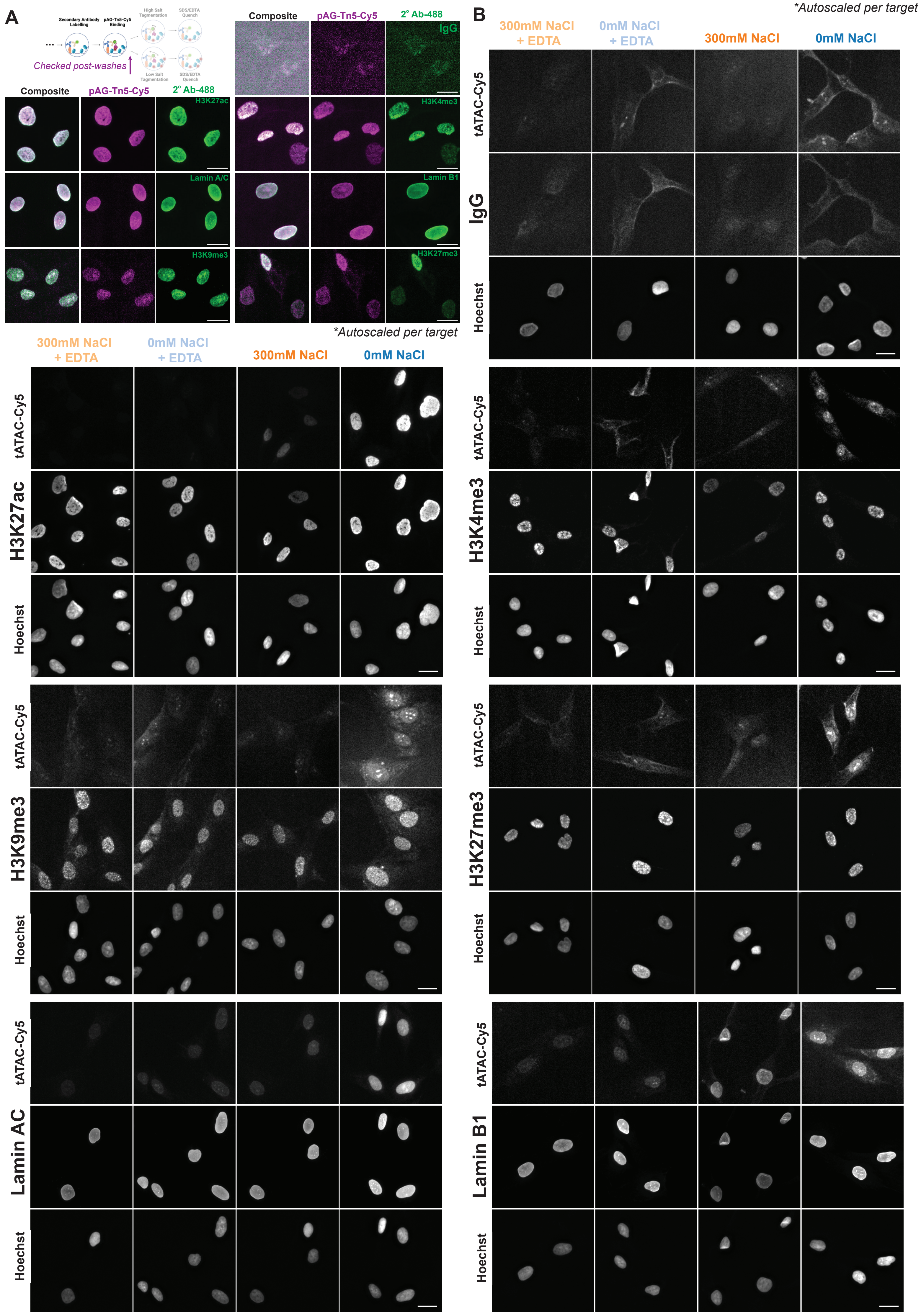
A
Secondary Antibody Labelling
pAG-Tn5-Cy5 Binding
High Salt Tagmentation
SDS/EDTA Quench
Low Salt Tagmentation
Checked post-washes
Composite
pAG-Tn5-Cy5
2° Ab-488
IgG
H3K27ac
H3K4me3
Lamin A/C
Lamin B1
H3K9me3
H3K27me3
*Autoscaled per target
300mM NaCl + EDTA
0mM NaCl + EDTA
300mM NaCl
0mM NaCl
tATAC-Cy5
H3K27ac
Hoechst
H3K9me3
Lamin AC
B
*Autoscaled per target
IgG
H3K4me3
H3K27me3
Lamin B1

**Supplemental Figure 1. Visual validation of tATAC-see specificity.** (A) Representative images captured post pAG-Tn5-Cy5 incubation and high-salt washes, but prior to tagmentation, to demonstrate specific localization of pAG-Tn5-Cy5 to the target, and sufficient removal of non-specific DNA binding in IgG controls. (B) Representative immunofluorescence panels of active tATAC-see reactions (0 mM and 300 mM NaCl) adjacent to their respective EDTA-inhibited controls (+ EDTA). Across targeted histone modifications and architectural proteins, the addition of EDTA diminishes the tATAC-Cy5 signal, visually confirming that fluorescence is driven by Tn5 tagmentation rather than non-specific accumulation. Images captured using high-throughput widefield microscopy. Scale bars = 20 μm.

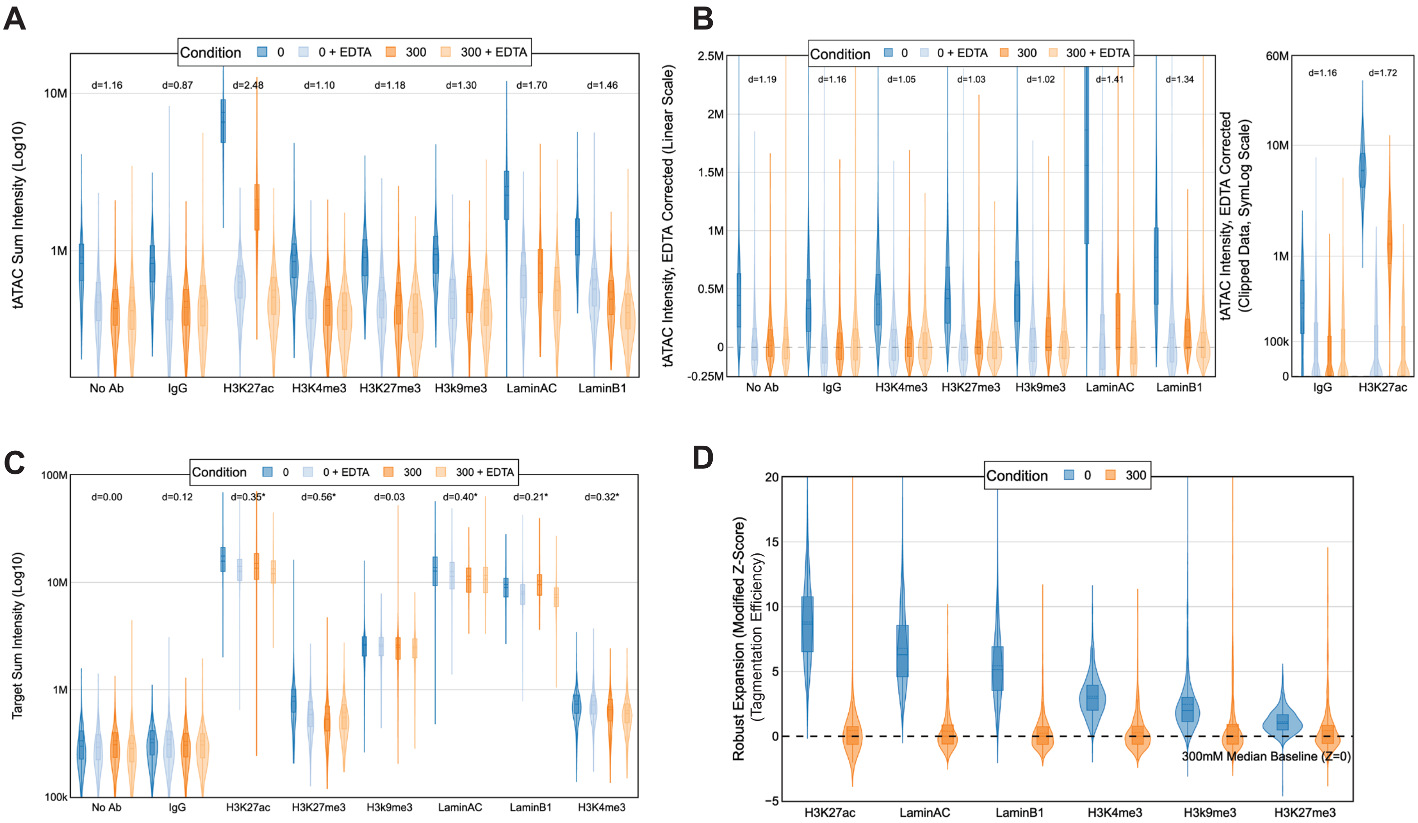

A
Condition
0
0 + EDTA
300
300 + EDTA
tATAC Sum Intensity (Log10)
10M
1M
d=1.16
d=0.87
d=2.48
d=1.10
d=1.18
d=1.30
d=1.70
d=1.46
No Ab
IgG
H3K27ac
H3K4me3
H3K27me3
H3k9me3
LaminAC
LaminB1
B
tATAC Intensity, EDTA Corrected (Linear Scale)
2.5M
2M
1.5M
1M
0.5M
0
-0.25M
d=1.19
d=1.16
d=1.05
d=1.03
d=1.02
d=1.41
d=1.34
No Ab
IgG
H3K4me3
H3K27me3
H3k9me3
LaminAC
LaminB1
tATAC Intensity, EDTA Corrected (Clipped Data, SymLog Scale)
60M
10M
1M
100k
0
d=1.16
d=1.72
IgG
H3K27ac
C
Target Sum Intensity (Log10)
100M
10M
1M
100k
d=0.00
d=0.12
d=0.35*
d=0.56*
d=0.03
d=0.40*
d=0.21*
d=0.32*
No Ab
IgG
H3K27ac
H3K27me3
H3k9me3
LaminAC
LaminB1
H3K4me3
D
Robust Expansion (Modified Z-Score) (Tagmentation Efficiency)
20
15
10
5
0
-5
300mM Median Baseline (Z=0)
H3K27ac
LaminAC
LaminB1
H3K4me3
H3k9me3
H3K27me3

**Supplemental Figure 2. Quantitative validation of tATAC-see via high-throughput widefield microscopy imaging.** (A) Violin plots of Raw tATAC Sum Intensity (Log10 scale) prior to EDTA (background) subtraction. (B) Violin plots displaying the tATAC signal (EDTA-Corrected) on a linear scale for lower intensity targets and a SymLog Scale using values clipped at 0 for H3k27ac, a high intensity target. Active single-cell values were corrected by subtracting the median ATAC signal of the corresponding target-matched EDTA control to isolate pure enzymatic activity. Statistics were conducted on unclipped values. (C) Violin plots of Target Sum Intensity (Log10 scale) demonstrating that target protein abundance minimally fluctuates across salt concentrations and EDTA treatments within each antibody group, though some statistical differences are noted. (D) Robust expansion of tagmentation efficiency (EDTA-Corrected), represented as a Modified Z-Score calculated against the 300 mM median baseline (Z=0 dashed line). Violin plots illustrate the kernel density distribution of single cells. Inner box plots denote the median and IQR. To quantify the magnitude of signal shifts independent of sample size, Cohen's *d* effect sizes comparing the 0 mM and 300 mM active conditions are denoted above each respective group in panels A–C. (A-D) Data shown are representative of a single 96-well plate experiment, which was performed twice for the full salt titration and more than three times for 0 and 300 mM pairs (*N* = 2 ; > 3 biological replicates). A minimum of *n* > 800 single nuclei were analyzed per condition. For A,B, all 0 mM vs 300 mM comparisons yielded p < 0.0001 via two-sided Mann-Whitney U test. For C, No Ab is not significant; IgG and H3K9me3, **p < 0.01; Lamin B1, ***p < 0.001; else ***p < 0.0001.

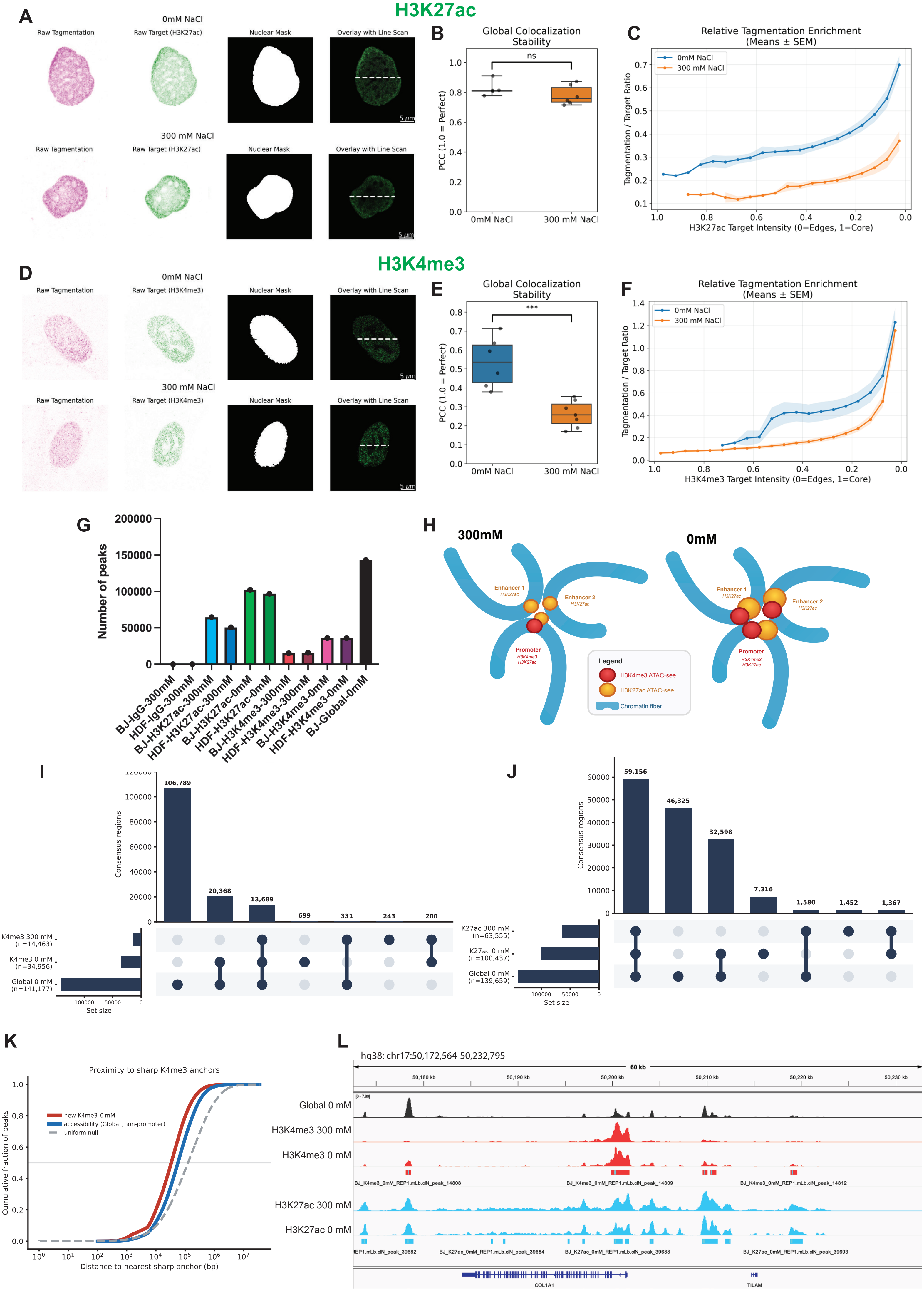

H3K27ac
A
0mM NaCl
Raw Tagmentation
Raw Target (H3K27ac)
Nuclear Mask
Overlay with Line Scan
300 mM NaCl
B
Global Colocalization Stability
ns
PCC (1.0 = Perfect)
0mM NaCl
300 mM NaCl
C
Relative Tagmentation Enrichment (Means ± SEM)
Tagmentation / Target Ratio
H3K27ac Target Intensity (0=Edges, 1=Core)
H3K4me3
D
Raw Target (H3K4me3)
E
***
F
H3K4me3 Target Intensity (0=Edges, 1=Core)
G
Number of peaks
BJ-IgG-300mM
HDF-IgG-300mM
BJ-H3K27ac-300mM
HDF-H3K27ac-300mM
BJ-H3K27ac-0mM
HDF-H3K27ac-0mM
BJ-H3K4me3-300mM
HDF-H3K4me3-300mM
BJ-H3K4me3-0mM
HDF-H3K4me3-0mM
BJ-Global-0mM
H
300mM
0mM
Enhancer 1
Enhancer 2
Promoter
Legend
H3K4me3 ATAC-see
H3K27ac ATAC-see
Chromatin fiber
I
Consensus regions
106,789
20,368
13,689
699
331
243
200
K4me3 300 mM (n=14,463)
K4me3 0 mM (n=34,956)
Global 0 mM (n=141,177)
Set size
J
59,156
46,325
32,598
7,316
1,580
1,452
1,367
K27ac 300 mM (n=63,555)
K27ac 0 mM (n=100,437)
Global 0 mM (n=139,659)
K
Proximity to sharp K4me3 anchors
new K4me3 0 mM
accessibility (Global, non-promoter)
uniform null
Cumulative fraction of peaks
Distance to nearest sharp anchor (bp)
L
hg38: chr17:50,172,564-50,232,795
60 kb
Global 0 mM
H3K4me3 300 mM
H3K4me3 0 mM
H3K27ac 300 mM
H3K27ac 0 mM
COL1A1
TILAM

**Supplemental Figure 3: High-resolution mapping of tATAC-see demonstrates target-proximal accessible microenvironment of active histone modifications.** (A-F) Airyscan SR imaging quantification for H3K27ac (A-C) and H3K4me3 (D-F) in neonatal fibroblasts. (A, D) Representative image analysis workflow. Each panel displays the isolated raw tagmentation signal (magenta), target chromatin signal (green), the algorithmically generated nuclear exclusion mask, and the merged spatial overlay. Dashed lines indicate the region used for the representative line scans shown in Fig. 2B, E. Scale bars = 5 µm. (B, E) Global colocalization stability between the tagmentation and target channels, measured via Pearson Correlation Coefficient (PCC) of masked nuclear pixels. Boxplot whiskers represent the 0th and 100th percentiles to display the full biological variance, with raw single-cell data points overlaid. (C, F) Relative tagmentation enrichment profiled across the target density gradient. Normalized target chromatin pixels were binned by target intensity from the low-intensity domain edges (0.0) to the high-intensity core (1.0). Curves represent the mean tagmentation-to-target ratio per bin across all analyzed cells. Shaded error bands represent the Standard Error of the Mean (SEM). Independent Student's t-tests were performed on $n$ = 5; 6; 6; 7 single nuclei per condition for H3K27ac 0 and 300 mM, and H3K4me3 0 mM and 300 mM, respectively, (*$p < 0.05$, ***$p < 0.001$). (G-L) Genomic sequencing following the tATAC-see protocol performed on two independent cell lines. (G) Number of MACS2 narrowpeaks called on IgG, H3K27ac, H3K4me3 (each at 300 mM and 0 mM NaCl) in neonatal and 22-year-old HDFs, alongside the neonatal Global ATAC-see. IgG controls yielded negligible peaks in both cell types, and peak numbers were concordant between neonatal and 22-year-old HDFs. (H) Schematic of the salt-titration spatial principle. Active regulatory elements brought together in a promoter–enhancer hub are shown on radiating chromatin fibers: a promoter marked by H3K4me3 and H3K27ac, and two H3K27ac-marked enhancers. At high ionic strength (300 mM), antibody-targeted tagmentation resolves sharp, focal signal that marks each element strictly at its own position. At low ionic strength (0 mM), recovery extends beyond the directly bound nucleosomes into the surrounding physically associated chromatin, capturing additional signal across the interacting hub. This broadening of focal peaks into interaction-hub detection at 0 mM is more pronounced for H3K4me3 than for H3K27ac. The model explains the underlying observations in Fig. 2G-L. Red, H3K4me3 tATAC-see signal; orange, H3K27ac tATAC-see signal; blue, chromatin fiber. (I, J) UpSet plots of peak-set overlap for H3K4me3 (I) and H3K27ac (J) across the same-mark 0 mM, 300 mM, and Global 0 mM peak sets on the standard chromosomes. (I) H3K4me3 (universe = 142,319 regions from 193,789 peaks) is dominated by the Global-only fraction (106,789; 75.0%), followed by H3K4me3 0 mM ∩ Global (20,368; 14.3%) and the all-three intersection (13,689; 9.6%), with H3K4me3-specific classes each ≤0.5%. Approximately 97% of H3K4me3 regions (both salt conditions) overlapped a Global peak. (J) H3K27ac (universe = 149,794 regions from 309,603 peaks) is dominated by the all-three intersection (59,156; 39.5%), followed by Global-only (46,325; 30.9%) and H3K27ac 0 mM ∩ Global (32,598; 21.8%), with H3K27ac-only and other minor classes each ≤4.9%. ≥91% of H3K27ac regions overlapped a Global peak. (K) Empirical cumulative distribution of the distance from each gained H3K4me3 0 mM peak (peaks not overlapping any 300 mM peak; $n$ = 21,524) to the nearest sharp 300 mM H3K4me3 summit (red), compared against the same distance computed for the non-promoter accessibility background (Global ATAC-see peaks not overlapping a 300 mM peak; $n$ = 127,659; blue) and a uniform null where gained-peak summits were randomized within each chromosome's peak span (grey dashed, 5 randomizations pooled). Gained peaks were significantly closer to sharp anchors than the accessibility background (median 33.3 kb vs 56.6 kb; uniform null 125.5 kb; one-sided Mann–Whitney $p \approx 0$, Cliff's $\delta = +0.21$). All distances are summit-to-summit and computed within chromosomes. (L) Representative IGV track view (hg38, chr17:50,172,564–50,232,795; ~60 kb) at the *COL1A1* locus showing Global ATAC-see, H3K4me3 300 mM and 0 mM, and H3K27ac 300 mM and 0 mM, with the corresponding 0 mM MACS2 peak calls below each mark.

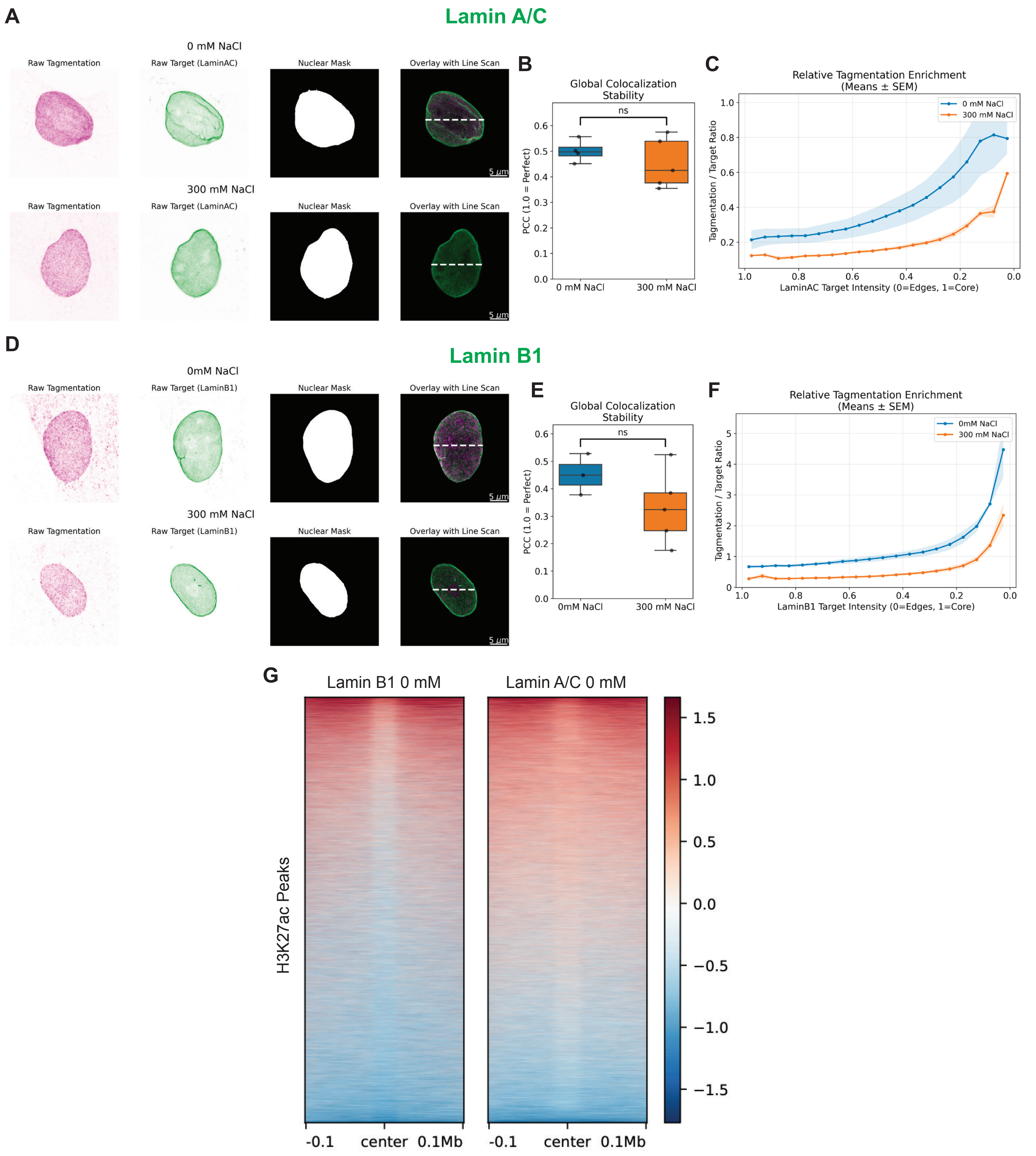
A
Lamin A/C
0 mM NaCl
Raw Tagmentation
Raw Target (LaminAC)
Nuclear Mask
Overlay with Line Scan
5 µm
300 mM NaCl
Raw Tagmentation
Raw Target (LaminAC)
Nuclear Mask
Overlay with Line Scan
5 µm
B
Global Colocalization Stability
ns
PCC (1.0 = Perfect)
0 mM NaCl
300 mM NaCl
C
Relative Tagmentation Enrichment (Means ± SEM)
0 mM NaCl
300 mM NaCl
Tagmentation / Target Ratio
LaminAC Target Intensity (0=Edges, 1=Core)
D
Lamin B1
0mM NaCl
Raw Tagmentation
Raw Target (LaminB1)
Nuclear Mask
Overlay with Line Scan
5 µm
300 mM NaCl
Raw Tagmentation
Raw Target (LaminB1)
Nuclear Mask
Overlay with Line Scan
5 µm
E
Global Colocalization Stability
ns
PCC (1.0 = Perfect)
0mM NaCl
300 mM NaCl
F
Relative Tagmentation Enrichment (Means ± SEM)
0mM NaCl
300 mM NaCl
Tagmentation / Target Ratio
LaminB1 Target Intensity (0=Edges, 1=Core)
G
Lamin B1 0 mM
Lamin A/C 0 mM
H3K27ac Peaks
-0.1
center
0.1Mb
-0.1
center
0.1Mb

**Supplemental Figure 4: High-resolution mapping of tATAC-see demonstrates target-proximal tagmentable DNA at the nuclear lamina.** (A-F) Airyscan SR imaging quantification for Lamin A/C (A-C) and Lamin B1 (D-F) in neonatal fibroblasts. (A, D) Representative image analysis workflow. Each panel displays the isolated raw tagmentation signal (magenta), target chromatin signal (green), the algorithmically generated nuclear exclusion mask, and the merged spatial overlay. Dashed lines indicate the regions used for the representative line scans shown in Fig. 3B, E. Scale bars = 5 µm. (B, E) Global colocalization stability between the tagmentation and target channels, measured via Pearson Correlation Coefficient (PCC) of masked nuclear pixels. Boxplot whiskers represent the 0th and 100th percentiles to display the full biological variance, with raw single-cell data points overlaid. (C, F) Relative tagmentation enrichment profiled across the target density gradient. Normalized target chromatin pixels were binned by target intensity from the low-intensity domain edges (0.0) to the high-intensity core (1.0). Curves represent the mean tagmentation-to-target ratio per bin across all analyzed cells. Shaded error bands represent the Standard Error of the Mean (SEM). Independent Student's t-tests were performed on $n$ = 4; 5; 4; 5 single nuclei per condition for Lamin A/C 0 and 300 mM, and Lamin B1 0 mM and 300 mM, respectively (ns = not significant). (G) Heatmaps of IgG normalized Lamin B1 (left) and Lamin A/C (right) targeted 0 mM ATAC-see signal across a ±100-kb window centered on H3K27ac targeted ATAC-see peak centres. A reduction in lamin signal centred around the H3K27ac peak is consistent with active regulatory elements being locally excluded from peripheral lamina contact, and the exclusion is more pronounced for Lamin B1 than Lamin A/C. Observations were confirmed for 2 independent cell lines.

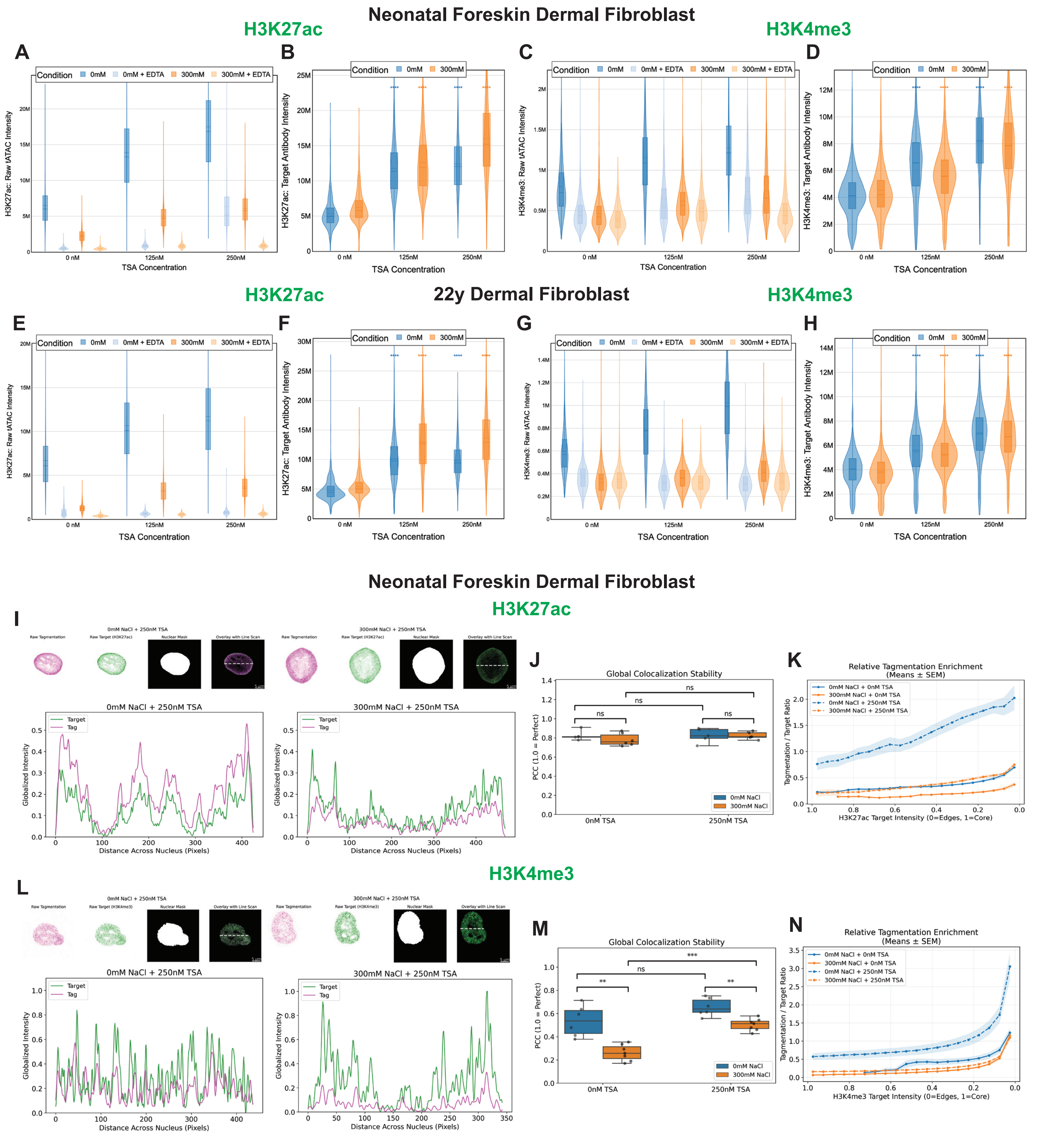

Neonatal Foreskin Dermal Fibroblast
H3K27ac
H3K4me3
A
B
C
D
Condition
0mM
0mM + EDTA
300mM
300mM + EDTA
H3K27ac: Raw tATAC Intensity
H3K27ac: Target Antibody Intensity
H3K4me3: Raw tATAC Intensity
H3K4me3: Target Antibody Intensity
TSA Concentration
0 nM
125nM
250nM
22y Dermal Fibroblast
E
F
G
H
Neonatal Foreskin Dermal Fibroblast
H3K27ac
I
0mM NaCl + 250nM TSA
300mM NaCl + 250nM TSA
Raw Tagmentation
Raw Target (H3K27ac)
Nuclear Mask
Overlay with Line Scan
Target
Tag
Globalized Intensity
Distance Across Nucleus (Pixels)
J
Global Colocalization Stability
PCC (1.0 = Perfect)
ns
0nM TSA
250nM TSA
0mM NaCl
300mM NaCl
K
Relative Tagmentation Enrichment
(Means ± SEM)
0mM NaCl + 0nM TSA
300mM NaCl + 0nM TSA
0mM NaCl + 250nM TSA
300mM NaCl + 250nM TSA
Tagmentation / Target Ratio
H3K27ac Target Intensity (0=Edges, 1=Core)
H3K4me3
L
Raw Target (H3K4me3)
M
**
***
N
H3K4me3 Target Intensity (0=Edges, 1=Core)

**Supplemental Figure 5: Expansion of euchromatic domains following TSA treatment.** (A-H) Quantitative evaluation of single-cell imaging intensities for neonatal (A–D) and 22-year-old (E–H) HDFs for H3K27ac (A, B, E, F) and H3K4me3 (C, D, G, H) following 4-hour TSA treatment at 0, 125, and 250 nM. (A, C, E, G) Raw tATAC-see intensities, including EDTA-inhibited controls. (B, D, F, H) Target antibody intensities across TSA treatments. All violin plots represent single-cell distributions; box plot overlays indicate medians and interquartile ranges. (I-N) Airyscan SR imaging quantification for H3K27ac (I-K) and H3K4me3 (L-N) in neonatal fibroblasts following 4-hour TSA treatment at 250 nM. (I, L) Upper panels: Representative image analysis workflow. Each panel displays the isolated raw tagmentation signal (magenta), target chromatin signal (green), the algorithmically generated nuclear exclusion mask, and the merged spatial overlay. Dashed lines indicate the regions used for the representative line scans below. Scale bars = 5 μm. Lower panels: Representative spatial line scans mapping the globally-scaled fluorescence intensities of the target (green) and tagmentation (magenta) channels at the cross-section of the nucleus marked above. Baseline 0 nM TSA control data are presented in Fig. 2B, E. (J, M) Global colocalization stability between the tagmentation and target channels, measured via Pearson Correlation Coefficient (PCC) of masked nuclear pixels. Boxplot whiskers represent the 0th and 100th percentiles to display the full biological variance, with raw single-cell data points overlaid. (K, N) Relative tagmentation enrichment profiled across the target density gradient. Normalized target chromatin pixels were binned by target intensity from the low-intensity domain edges (0.0) to the high-intensity core (1.0). Curves represent the mean tagmentation-to-target ratio per bin across all analyzed cells. Shaded error bands represent the Standard Error of the Mean (SEM). For widefield imaging data (A-H), significance was determined via two-sided Mann-Whitney U tests on unclipped data for $n \geq 420$ single nuclei per condition (****$p < 0.0001$). Data is a representative single 96-well plate experiment of more than $N \geq 2$ biological replicates. For SR imaging (J, M), statistical significance was determined via independent Student's t-tests for $n = 5$ or 6 single nuclei per condition (**$p < 0.01$, ***$p < 0.001$, ns = not significant).

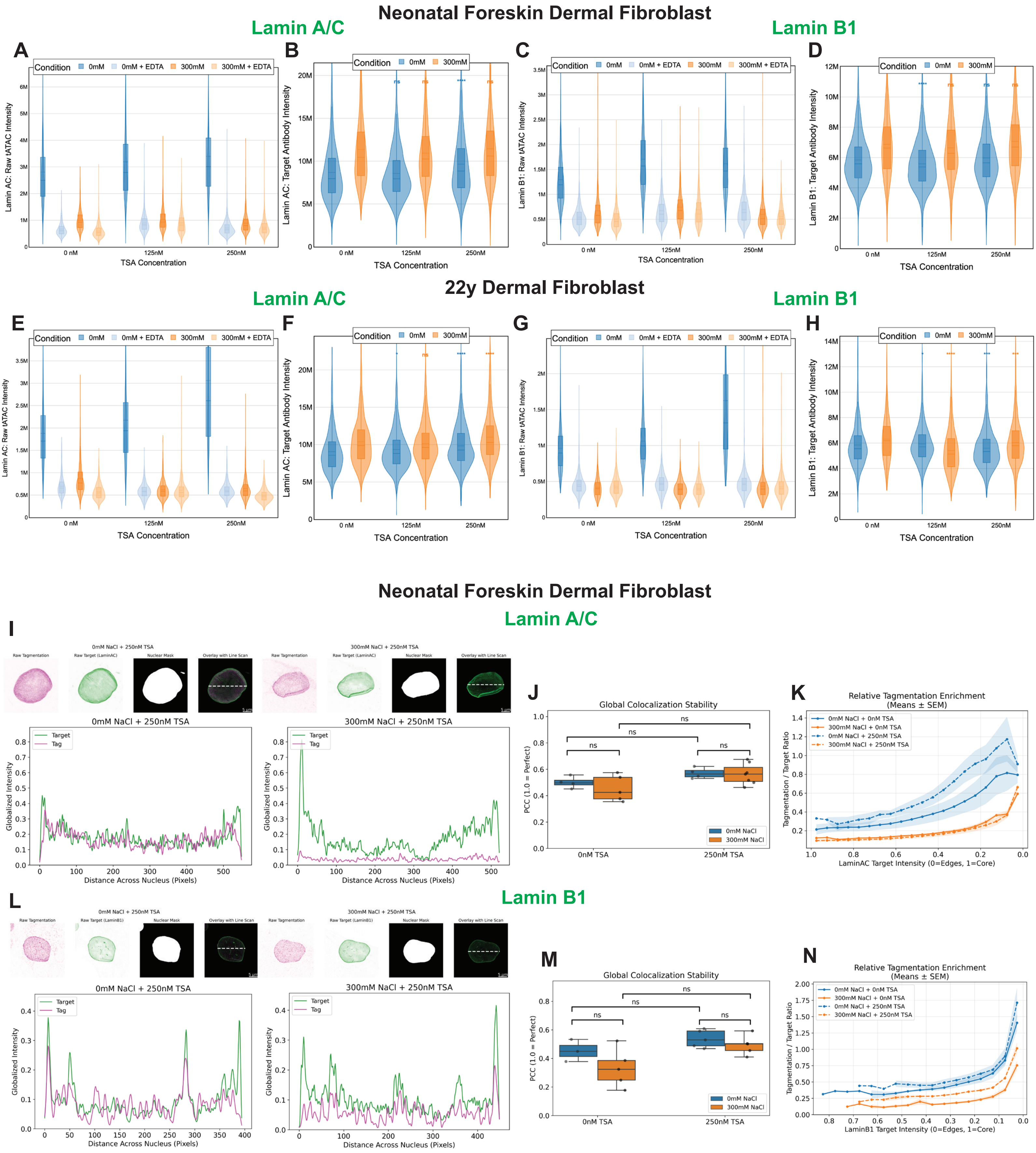

Neonatal Foreskin Dermal Fibroblast
Lamin A/C
Lamin B1
A
Condition 0mM 0mM + EDTA 300mM 300mM + EDTA
Lamin AC: Raw tATAC Intensity
TSA Concentration
0 nM
125nM
250nM
B
Condition 0mM 300mM
Lamin AC: Target Antibody Intensity
TSA Concentration
C
Condition 0mM 0mM + EDTA 300mM 300mM + EDTA
Lamin B1: Raw tATAC Intensity
TSA Concentration
D
Condition 0mM 300mM
Lamin B1: Target Antibody Intensity
TSA Concentration
22y Dermal Fibroblast
Lamin A/C
Lamin B1
E
Condition 0mM 0mM + EDTA 300mM 300mM + EDTA
Lamin AC: Raw tATAC Intensity
TSA Concentration
F
Condition 0mM 300mM
Lamin AC: Target Antibody Intensity
TSA Concentration
G
Condition 0mM 0mM + EDTA 300mM 300mM + EDTA
Lamin B1: Raw tATAC Intensity
TSA Concentration
H
Condition 0mM 300mM
Lamin B1: Target Antibody Intensity
TSA Concentration
Neonatal Foreskin Dermal Fibroblast
Lamin A/C
I
0mM NaCl + 250nM TSA
300mM NaCl + 250nM TSA
Raw Tagmentation
Raw Target (LaminAC)
Nuclear Mask
Overlay with Line Scan
0mM NaCl + 250nM TSA
300mM NaCl + 250nM TSA
Target
Tag
Globalized Intensity
Distance Across Nucleus (Pixels)
J
Global Colocalization Stability
PCC (1.0 = Perfect)
ns
0mM NaCl
300mM NaCl
0nM TSA
250nM TSA
K
Relative Tagmentation Enrichment
(Means ± SEM)
0mM NaCl + 0nM TSA
300mM NaCl + 0nM TSA
0mM NaCl + 250nM TSA
300mM NaCl + 250nM TSA
Tagmentation / Target Ratio
LaminAC Target Intensity (0=Edges, 1=Core)
Lamin B1
L
0mM NaCl + 250nM TSA
300mM NaCl + 250nM TSA
Raw Tagmentation
Raw Target (LaminB1)
Nuclear Mask
Overlay with Line Scan
0mM NaCl + 250nM TSA
300mM NaCl + 250nM TSA
Target
Tag
Globalized Intensity
Distance Across Nucleus (Pixels)
M
Global Colocalization Stability
PCC (1.0 = Perfect)
ns
0mM NaCl
300mM NaCl
0nM TSA
250nM TSA
N
Relative Tagmentation Enrichment
(Means ± SEM)
0mM NaCl + 0nM TSA
300mM NaCl + 0nM TSA
0mM NaCl + 250nM TSA
300mM NaCl + 250nM TSA
Tagmentation / Target Ratio
LaminB1 Target Intensity (0=Edges, 1=Core)

**Supplemental Figure 6: Remodeling of Lamina-Associated Domains following TSA treatment.** (A-H) Quantitative evaluation of single-cell imaging intensities for neonatal (A–D) and 22-year-old (E–H) HDFs for Lamin A/C (A, B, E, F) and Lamin B1 (C, D, G, H) following 4-hour TSA treatment at 0, 125, and 250 nM. (A, C, E, G) Raw tATAC-see intensities, including EDTA-inhibited controls. (B, D, F, H) Target antibody intensities across TSA treatments. All violin plots represent single-cell distributions; box plot overlays indicate medians and interquartile ranges. (I-N) Airyscan SR imaging quantification for Lamin A/C (I-K) and Lamin B1 (L-N) in neonatal fibroblasts following 4-hour TSA treatment at 250 nM. (I, L) Upper panels: Representative image analysis workflow. Each panel displays the isolated raw tagmentation signal (magenta), target chromatin signal (green), the algorithmically generated nuclear exclusion mask, and the merged spatial overlay. Dashed lines indicate the regions used for the representative line scans below. Scale bars = 5 µm. Lower panels: Representative spatial line scans mapping the globally-scaled fluorescence intensities of the target (green) and tagmentation (magenta) channels at the cross-section of the nucleus marked above. Baseline 0 nM TSA control data are presented in Fig. 3B, E. (J, M) Global colocalization stability between the tagmentation and target channels, measured via Pearson Correlation Coefficient (PCC) of masked nuclear pixels. Boxplot whiskers represent the 0th and 100th percentiles to display the full biological variance, with raw single-cell data points overlaid. (K, N) Relative tagmentation enrichment profiled across the target density gradient. Normalized target chromatin pixels were binned by target intensity from the low-intensity domain edges (0.0) to the high-intensity core (1.0). Curves represent the mean tagmentation-to-target ratio per bin across all analyzed cells. Shaded error bands represent the Standard Error of the Mean (SEM). For widefield imaging data (A-H), significance was determined via two-sided Mann-Whitney U tests on unclipped data for $n \geq 420$ nuclei per condition and is representative more than $N \geq 2$ biological replicates (*$p < 0.05$, **$p < 0.01$, ***$p < 0.001$, ****$p < 0.0001$, ns = not significant). For SR imaging (J, M), statistical significance was determined via independent Student's t-tests for $n$ = 4, 5 or 7 single nuclei per condition (ns = not significant).

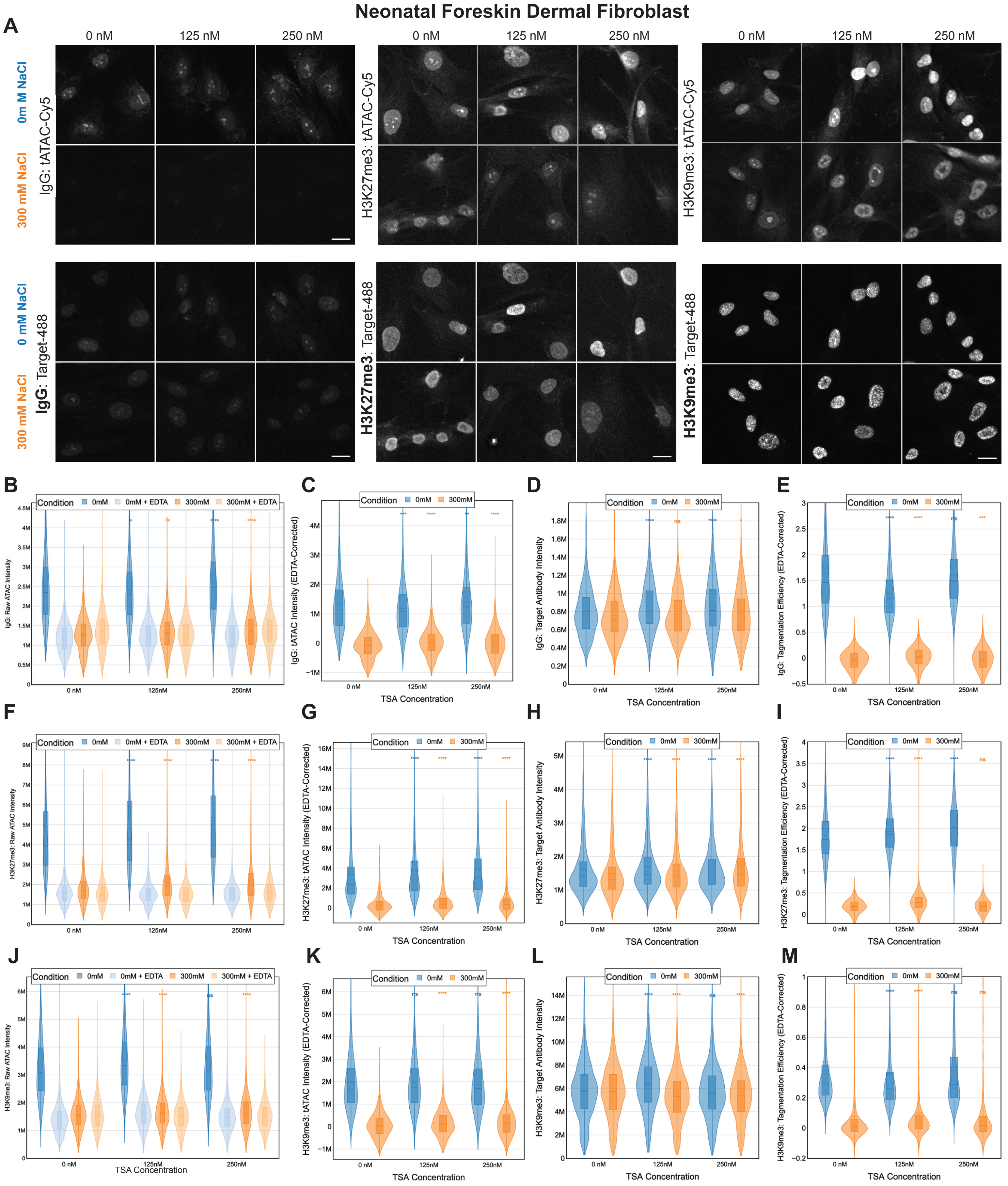
Neonatal Foreskin Dermal Fibroblast
A
0 nM
125 nM
250 nM
0m M NaCl
300 mM NaCl
IgG: tATAC-Cy5
H3K27me3: tATAC-Cy5
H3K9me3: tATAC-Cy5
0 mM NaCl
300 mM NaCl
IgG: Target-488
H3K27me3: Target-488
H3K9me3: Target-488
B
C
D
E
F
G
H
I
J
K
L
M
Condition
0mM
0mM + EDTA
300mM
300mM + EDTA
IgG: Raw ATAC Intensity
IgG: tATAC Intensity (EDTA-Corrected)
IgG: Target Antibody Intensity
IgG: Tagmentation Efficiency (EDTA-Corrected)
H3K27me3: Raw ATAC Intensity
H3K27me3: tATAC Intensity (EDTA-Corrected)
H3K27me3: Target Antibody Intensity
H3K27me3: Tagmentation Efficiency (EDTA-Corrected)
H3K9me3: Raw ATAC Intensity
H3K9me3: tATAC Intensity (EDTA-Corrected)
H3K9me3: Target Antibody Intensity
H3K9me3: Tagmentation Efficiency (EDTA-Corrected)
TSA Concentration
0 nM
125nM
250nM
ns

**Supplemental Figure 7: Quantitative widefield imaging of repressive histone marks and IgG controls following TSA treatment.** (A) Representative extended depth of focus (EDF) projections from widefield imaging of the tagmentation signal (tATAC-Cy5) and the structural anchor (Target-488) for IgG, H3K27me3, and H3K9me3 across TSA concentrations and salt conditions (0 mM vs. 300 mM NaCl). Scale bars = 20 µm. Note that tATAC-see signal for H3K27me3 and H3K9me3 is prominently distributed within nucleoli. (B-J) Quantitative evaluation of single-cell imaging intensities for neonatal HDFs following 4-hour TSA treatment at 0, 125, and 250 nM for IgG controls (B-D), H3K27me3 (E-G), and H3K9me3 (H-J). (B, F, J) Raw tATAC-see intensities, including EDTA-inhibited controls. (C, G, K) EDTA-Corrected tATAC-see intensities. (D, H, L) Target antibody intensities across TSA treatments. (E, I, M) Normalized tagmentation efficiency (EDTA-Corrected). All violin plots represent single-cell distributions; box plot overlays indicate medians and interquartile ranges. Significance was determined via two-sided Mann-Whitney U tests on unclipped data for $n \geq 1563$ nuclei per condition and is representative of $N = 3$ biological replicates (*$p < 0.05$, **$p < 0.01$, ***$p < 0.001$, ****$p < 0.0001$, ns = not significant).

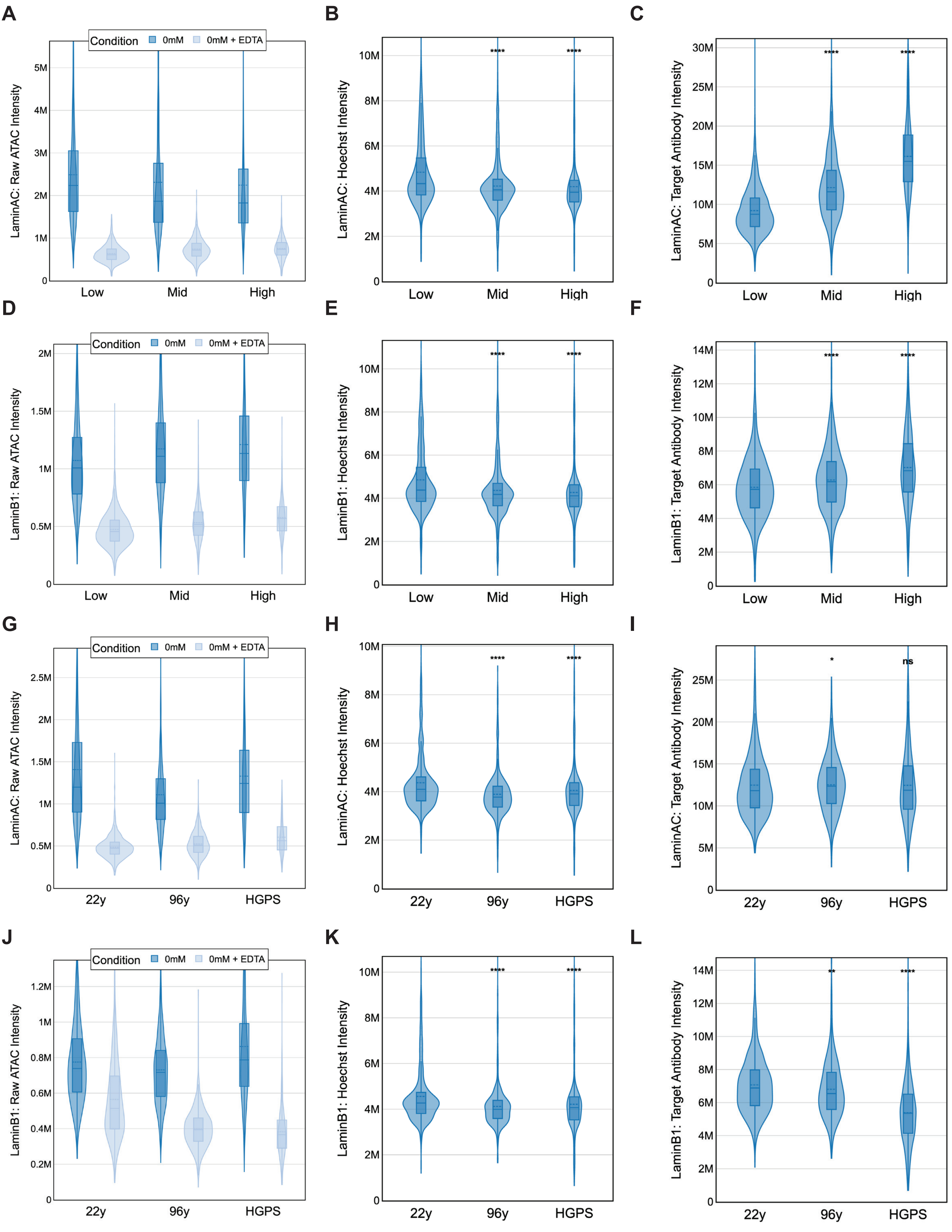
A
B
C
D
E
F
G
H
I
J
K
L
Condition
0mM
0mM + EDTA
LaminAC: Raw ATAC Intensity
LaminAC: Hoechst Intensity
LaminAC: Target Antibody Intensity
LaminB1: Raw ATAC Intensity
LaminB1: Hoechst Intensity
LaminB1: Target Antibody Intensity
Low
Mid
High
22y
96y
HGPS
****
**
*
ns

**Supplemental Figure 8: Quantitative validation of target abundance and technical baselines across aging models at 0 mM NaCl for Lamin A/C and B1.** (A-L) Quantitative evaluation of single-cell imaging intensities for Lamin A/C (A-C, G-I) and Lamin B1 (D-F, J-L) across models of replicative aging (Low, Mid, High) and chronological aging/HGPS (22y, 96y, HGPS). (A, D, G, J) Raw tATAC-see intensities for 0 mM NaCl reactions, including EDTA-inhibited controls. (B, E, H, K) Hoechst intensities. (C, F, I, L) Target antibody intensities. All violin plots represent single-cell distributions; box plot overlays indicate medians and interquartile ranges. Significance was determined via two-sided Mann-Whitney U tests on unclipped data for $n \geq 540$ single nuclei per condition and is representative of $N = 2$ biological replicates (*$p < 0.05$, **$p < 0.01$, ****$p < 0.0001$, ns = not significant).

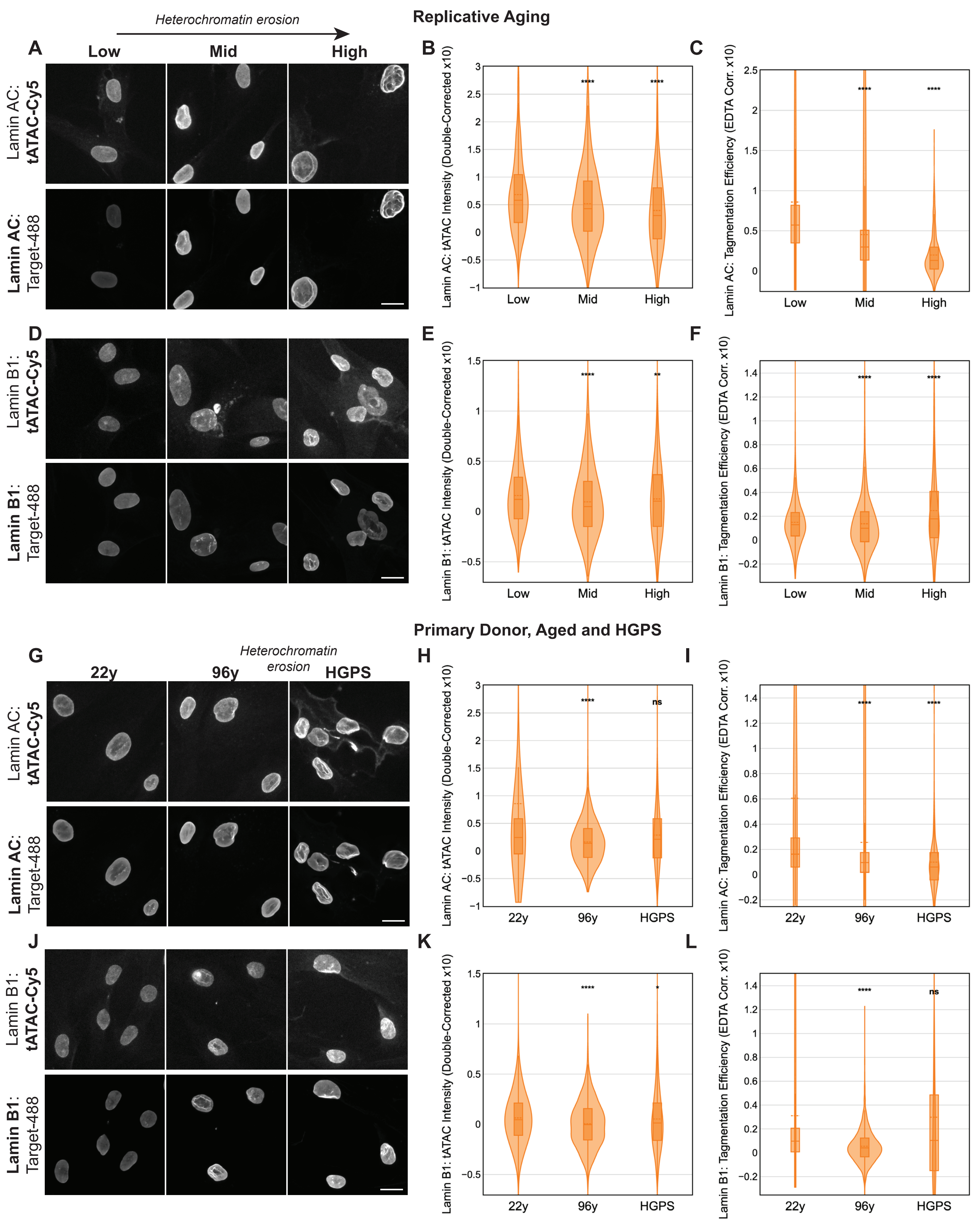
Replicative Aging
Heterochromatin erosion
A
Low
Mid
High
Lamin AC: tATAC-Cy5
Lamin AC: Target-488
B
Lamin AC: tATAC Intensity (Double-Corrected x10)
****
****
Low
Mid
High
C
Lamin AC: Tagmentation Efficiency (EDTA Corr. x10)
****
****
Low
Mid
High
D
Lamin B1: tATAC-Cy5
Lamin B1: Target-488
E
Lamin B1: tATAC Intensity (Double-Corrected x10)
****
**
Low
Mid
High
F
Lamin B1: Tagmentation Efficiency (EDTA Corr. x10)
****
****
Low
Mid
High
Primary Donor, Aged and HGPS
G
Heterochromatin erosion
22y
96y
HGPS
Lamin AC: tATAC-Cy5
Lamin AC: Target-488
H
Lamin AC: tATAC Intensity (Double-Corrected x10)
****
ns
22y
96y
HGPS
I
Lamin AC: Tagmentation Efficiency (EDTA Corr. x10)
****
****
22y
96y
HGPS
J
Lamin B1: tATAC-Cy5
Lamin B1: Target-488
K
Lamin B1: tATAC Intensity (Double-Corrected x10)
****
*
22y
96y
HGPS
L
Lamin B1: Tagmentation Efficiency (EDTA Corr. x10)
****
ns
22y
96y
HGPS

**Supplemental Figure 9: Quantitative widefield tATAC-see mapping of lamina-associated chromatin under high-stringency (300 mM NaCl) conditions.** (A, D, G, J) Representative extended depth of focus (EDF) projections from widefield imaging of the tagmentation signal (tATAC-Cy5) and the structural anchor (Target-488) for Lamin A/C (A, G) and Lamin B1 (D, J) at 300 mM NaCl. Images display replicative aging (Low, Mid, High) and chronological aging/HGPS (22y, 96y, HGPS) models. Scale bars = 20 μm. (B, E, H, K) Total single-cell tATAC-see intensity (Double-Corrected for DNA content and EDTA background) for Lamin A/C (B, H) and Lamin B1 (E, K). (C, F, I, L) Normalized tagmentation efficiency (EDTA-Corrected) for Lamin A/C (C, I) and Lamin B1 (F, L). All violin plots represent single-cell distributions; box plot overlays indicate medians and interquartile ranges. Significance was determined via two-sided Mann-Whitney U tests on unclipped data for $n \geq 500$ nuclei per condition (****$p < 0.0001$, ns = not significant).

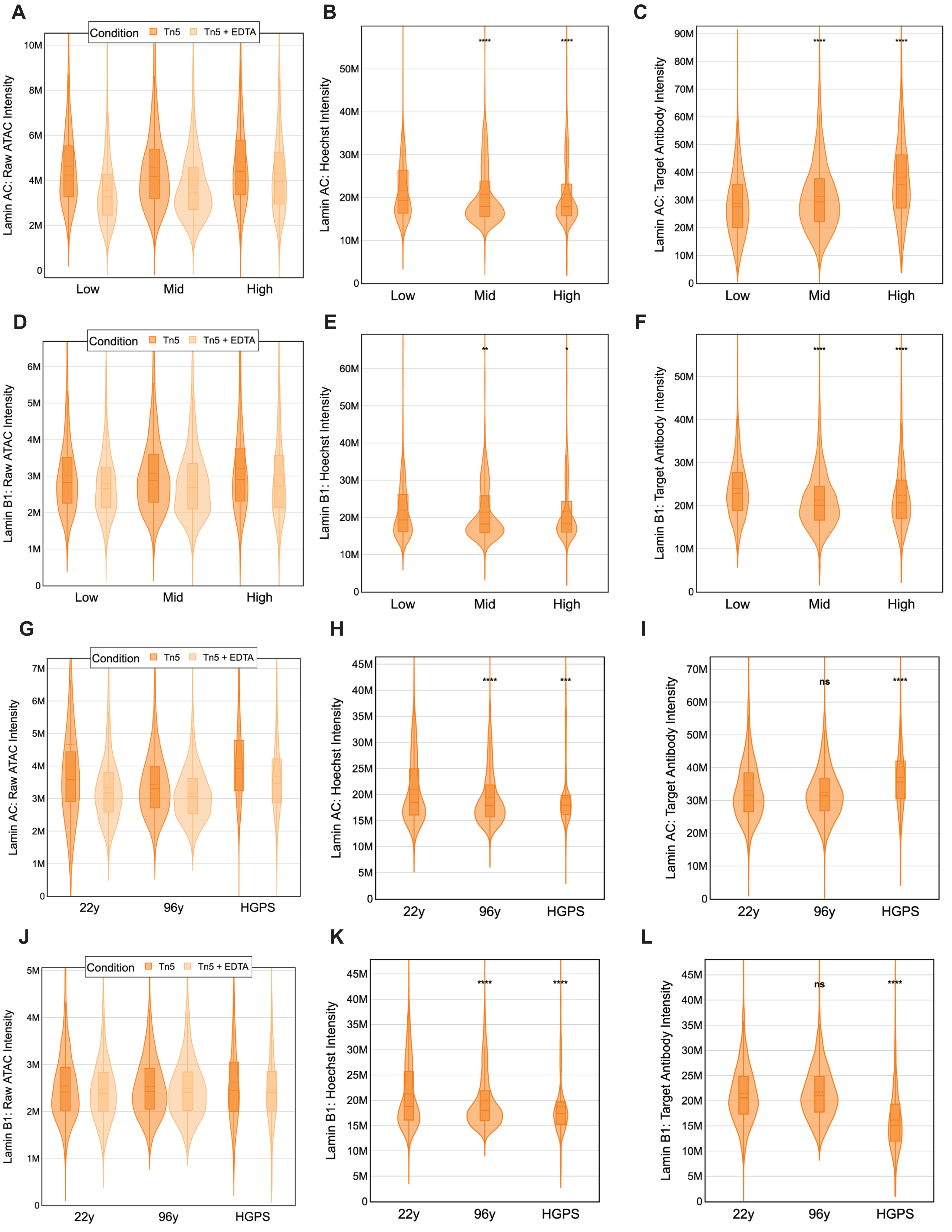

A
Condition
Tn5
Tn5 + EDTA
Lamin AC: Raw ATAC Intensity
0
2M
4M
6M
8M
10M
Low
Mid
High
B
Lamin AC: Hoechst Intensity
0
10M
20M
30M
40M
50M
****
****
Low
Mid
High
C
Lamin AC: Target Antibody Intensity
0
10M
20M
30M
40M
50M
60M
70M
80M
90M
****
****
Low
Mid
High
D
Condition
Tn5
Tn5 + EDTA
Lamin B1: Raw ATAC Intensity
0
1M
2M
3M
4M
5M
6M
Low
Mid
High
E
Lamin B1: Hoechst Intensity
0
10M
20M
30M
40M
50M
60M
**
*
Low
Mid
High
F
Lamin B1: Target Antibody Intensity
0
10M
20M
30M
40M
50M
****
****
Low
Mid
High
G
Condition
Tn5
Tn5 + EDTA
Lamin AC: Raw ATAC Intensity
0
1M
2M
3M
4M
5M
6M
7M
22y
96y
HGPS
H
Lamin AC: Hoechst Intensity
0
5M
10M
15M
20M
25M
30M
35M
40M
45M
****
***
22y
96y
HGPS
I
Lamin AC: Target Antibody Intensity
0
10M
20M
30M
40M
50M
60M
70M
ns
****
22y
96y
HGPS
J
Condition
Tn5
Tn5 + EDTA
Lamin B1: Raw ATAC Intensity
0
1M
2M
3M
4M
5M
22y
96y
HGPS
K
Lamin B1: Hoechst Intensity
0
5M
10M
15M
20M
25M
30M
35M
40M
45M
****
****
22y
96y
HGPS
L
Lamin B1: Target Antibody Intensity
0
5M
10M
15M
20M
25M
30M
35M
40M
45M
ns
****
22y
96y
HGPS

**Supplemental Figure 10: Quantitative validation of target abundance and technical baselines across aging models at 300 mM NaCl.** (A-L) Quantitative evaluation of single-cell imaging intensities for Lamin A/C (A-C, G-I) and Lamin B1 (D-F, J-L) across models of replicative aging (Low, Mid, High) and chronological aging/HGPS (22y, 96y, HGPS). (A, D, G, J) Raw tATAC-see intensities for 300 mM NaCl reactions (labeled Tn5 vs. Tn5 + EDTA). (B, E, H, K) Hoechst intensities. (C, F, I, L) Target antibody intensities. All violin plots represent single-cell distributions; box plot overlays indicate medians and interquartile ranges. Significance was determined via two-sided Mann-Whitney U tests on unclipped data for $n \geq 500$ nuclei per condition (****$p < 0.0001$, ns = not significant).
